# Annual Variability in Diffuse Ratio and Spectral Characteristics of Solar Radiation: Cloud Effects in a Temperate Monsoon Region

**DOI:** 10.1101/2024.10.14.618107

**Authors:** Amila Nuwan Siriwardana, Atsushi Kume

## Abstract

Solar radiation (SR) dynamics have a profound effect on plant growth, development, and ecosystems, and they act as a primary energy source and important environmental signal that plants perceive through their photoreceptors, which primarily sense critical wavelength ratios (CWRs). The diffuse fraction of solar radiation (DF) is a key factor affecting the quality and distribution of light within the plant canopy. We analyzed one year of SR spectral observations measured by a ground-based rotating shadow-band spectroradiometer to evaluate the potential effects of DF and CWRs on plants in an outdoor environment in Fukuoka, Japan. The daily mean DF and all considered CWRs showed significant seasonal variations regardless of the solar meridian altitude. Cloudy or partly cloudy skies were prevalent throughout the year. The ratios of ultraviolet-A (UV-A)/UV-B, red (R)/blue (B), and R/green (G) increased during winter and decreased during summer. Conversely, the ratios of photosynthetically active radiation (PAR)/global solar radiation (GSI), UV/GSI, UV/PAR, B/G, R/far-red (FR), and UV-B/B increased during summer and decreased during winter. Specific CWRs correlated with air mass (AM) and vapor pressure (VP), and most of the CWR correlated with DF. AM, VP, and DF were also found to have a combined effect on specific CWRs that are important for plant light signals. This highlights the potential influence of AM, VP, and DF on plant light signals, thereby opening avenues for the development of innovative models of plant growth and ecological responses that incorporate plant photoreception.

## Introduction

Solar radiation (SR), which is a broad spectrum of electromagnetic radiation emitted by the sun, profoundly influences the Earth’s climate and a variety of life forms (Blal et al. 2020; Lozano et al. 2022). Some of SR penetrates the Earth’s atmosphere and reaches the Earth’s surface, of which approximately 6% is ultraviolet radiation (UVR) (λ 200–400 nm), approximately 52% is visible/ photosynthetically active radiation (PAR) (λ 400–700 nm), and the remaining 42% is infrared radiation (IR) (λ 760–10^6^ nm) (Polefka et al. 2012). Together, this provides approximately 1,361 W m^−2^ of energy to the Earth as a primary energy source (Solanki et al. 2013) that controls numerous processes. SR drives the physical and biochemical processes of transpiration and photosynthesis in plants, which affect the growth and development of plants during their life cycle (Yadav et al. 2020), and it determines net ecosystem productivity (NEP) (Yuan et al. 2014; Li et al. 2015).

The SR that passes through the Earth’s atmosphere is reflected, scattered, and absorbed by dust particles, gas molecules, ozone, and water vapor (Blal et al. 2020). This results in the partitioning of SR into two types: diffuse solar radiation (DFR) and direct solar radiation (DIR). DFR represents the fraction of light that is scattered, reflected, and reenters the Earth from multiple directions. In contrast, the SR received directly at the Earth’s surface without scattering is referred to as DIR (Mazza et al. 1999). The cumulative total of these two components is the global solar radiation (GSI) (Xin et al. 2016; Durand et al. 2021; Wu et al. 2022), and the ratio between DFR and GSI is called the diffuse factor or diffuse ratio (DF) (Shibuya et al. 2018). DIR and DFR propagate differently within canopies, which affects light distribution within the plant canopy (Xin et al. 2016; Durand et al. 2021; Wu et al. 2022), and has a significant influence on canopy photosynthesis and NEP (Blal et al. 2020). However, only a limited number of studies have addressed the effects of SR partitioning (Yuan et al. 2014; Xin et al. 2016), and the seasonal variation in DF remains poorly understood. In addition, many plant physiologists and photovoltaic researchers have conducted their studies assuming clear-sky conditions (Engerer and Mills 2015; Matthews et al. 2020; Peratikou and Charalambides 2022), and have not considered the numerous regions that have a limited number of clear-sky days.

The survival of organisms depends on their ability to accurately sense and respond to their extracellular environment. Many organisms have evolved sophisticated photosensory systems that allow them to respond to changes in SR (Chen et al. 2004). However, most research efforts consider SR primarily as an energy source (Yadav et al. 2020) and often overlook its multifaceted role as both an energy source and a central environmental signal (Shinomura et al. 1994; Kong and Okajima 2016; Yadav et al. 2020; Yang et al. 2020). SR provides specific light signals that regulate plant development and metabolism through complex photomorphogenetic mechanisms which is very important to the survival of stationary life of plants. These mechanisms respond to the quality of the light signal even at very low intensities, which is determined by its wavelengths (Yadav et al. 2020). Specifically, certain critical wavelength ratios (CWRs) play a significant role in plant light signaling (Mao et al. 2005; Yang et al. 2020). Specifically, we have chosen to investigate CWRs such as, Red(R) (λ 620–680 nm)/ far-red (FR) (λ 700–750 nm) (Tan et al. 2022), R/ blue (B) (λ 450–495nm)(S. Wang et al. 2022), R/(G) (λ 500–530 nm) (Trojak et al. 2022), B/G (Sellaro et al. 2010a), UV-A (λ 315–400 nm)/ UV-B (λ 280–315 nm) (Neugart and Schreiner 2018), UV-B/B (Tissot and Ulm 2020), UV/PAR, and UV/GSI(Rai et al. 2021). These light signals were selected due to their potentially significant impact on plant and ecosystem dynamics. Hence, this research aims to fill the knowledge gap of the seasonal variation in DF and CWRs specifically, R/B, R/G, B/G, GSI, and R/FR in addition to UV ratios under all weather conditions.

The changing cloud conditions that are directly reflected by the DF not only affect the Photosynthetically active photon flux density (PPFD) but may also alter the spectral distribution of SR and change the CWRs that lead to plant light signaling. Hence, we further investigated the influence of DF on CWRs to broaden the knowledge of such natural phenomena. The ratio of the path length of SR through the atmosphere relative to the path length from sea level to the zenith is the air mass (AM) (Malitson 1968). This could be simplified as the path length of light through the atmosphere. When the light path through the atmosphere, different wavelengths of SR, scatter differently hence the AM influences spectral solar irradiance (SSI) (Borchi et al. 2011), thus CWRs. Therefore, this research investigated the seasonal variation of AM and its effects on CWRs. Additionally, vapor pressure (VP) which is the partial pressure of water vapor in the atmosphere (Earthdata. earthdata.nasa.gov, 2024) also might influence the CWRs as water vapor absorption of SR depends on the wavelengths (Orte et al. 2021). Therefore, atmospheric VP changes could affect the CWRs thus plant light signals and ecosystem.

Based on year-round observations of DFR and DIR using a high-precision shadow band spectroradiometer, we clarified the actual state of SR diffusion and seasonal changes in the SR spectrum. This study aimed to quantify the seasonal variation of CWRs and DF, and construct models that accurately represent CWRs variations. Furthermore, we sought to understand the effects of VP, AM, and DF on CWRs. The findings from this research improve our understanding of natural SR fluctuation. The results indicate the potential influence of AM, VP, and DF on CWRs, thus plant light signals. Consequently, avenues for the development of innovative models for plant growth and ecological responses that incorporate plant photoreception are opened.

## Materials and Methods

The spectral radiation data were collected at Kyushu University, Ito Campus, which is located on the Itoshima Peninsula in western Fukuoka Prefecture, Kyushu island, Japan (33.594 N, 130.214 E; 90 m asl.). The region has a warm temperate climate (Cfa) and is strongly influenced by the monsoon. The average annual temperature (T) is 17.3°C, and the average annual precipitation is 1686.9 mm (AMeDAS Fukuoka station, 33.560 N, 130.090 E, 5 m asl.). The average annual temperature in 2021 was 18.2°C (Sup. 1a), bit the temperature varies widely between seasons (relative difference (*rd*) = 118% and absolute difference (*ad*) = 311%). Although the variation in relative humidity (RH) does not show a seasonal pattern (Sup. 1b), the annual average RH is high (61.3%). The local air pressure is low in summer (1007 hPa in August) and high in winter (1021 hPa in December) (Sup. 1c). The atmospheric water vapor pressure (VP) was calculated using RH and the saturation water vapor pressure (VPs), which was calculated using the Tetens equation (1930, Eq. 1) with the air temperature (*T*) in degrees Celsius (°C):

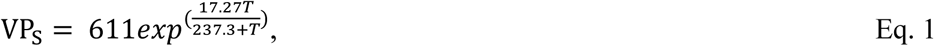

VP was calculated by multiplying VP_S_ by RH:

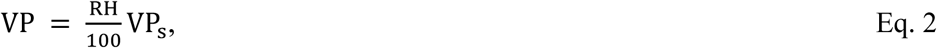

The VP and VP_S_ values increased during the summer and decreased during the winter season (Sup. 1d). The airmass (AM) is an important parameter that affects both the quality and quantity of light. According to Lambert’s cosine law (Eq. 3), the flux density at the surface changes with the zenith angle:

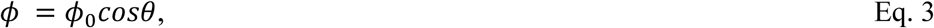

where *ϕ* is flux density at the surface, ϕ_0_ is the flux density normal to the beam and θ is the zenith angle. Moreover, the AM is change with the zenith angle according to Eq. 4:

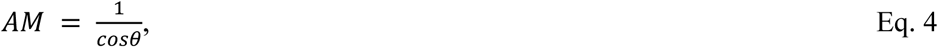

where AM is the airmass, and θ is the zenith angle. Furthermore, Bouguer’s (or Beer’s) law, which was developed from Lambert’s cosine law (Eq. 5), explains that the distance the beam travels reduces the flux density:

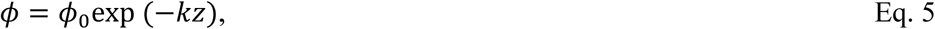

where ϕ_0_ is the flux density normal to the beam, *ϕ* is flux density at the sur, *k* is the extinction coefficient, and *z* is the distance the beam travels. Equation (Eq. 6) was derived from this equation to explain the importance of AM on the radiation intensity and to find the total atmospheric optical thickness (τ) (Borchi et al. 2011):

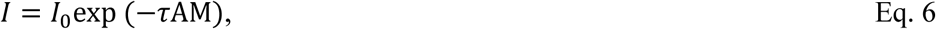

where *I* is the measured radiation intensity at ground level, and *I*_0_ is the intensity of radiation before it enters the atmosphere. Borchi et al. (2011) reported that the total atmospheric optical thickness (τ) varies with wavelength. The AM decreased during the summer and increased during the winter season (Sup. 1e). As AM increases, the peak of the SSI shifts toward longer wavelengths because shorter wavelengths are attenuated by Rayleigh scattering (Guechi et al. 2013). Perliski and Solomon (1993) recorded that the shorter visible wavelengths (<450 nm) scatter more at stratospheric AM than the longer wavelengths, whereas longer visible wavelengths (>650 nm) may scatter more at tropospheric AM than the shorter visible wavelengths. This indicates possible changes in CWRs and their influence on plant light signals. This study focused on the effect of AM on light signals in addition to VP and DF.

Radiometric measurements were taken using an EKO MS-711 spectroradiometer (S18056.06, EKO Instruments, Japan) and an EKO MB-22 rotating shadow band (RSB) (ES18239.01, EKO Instruments, Japan). The detector core of the EKO MS-711 is temperature controlled to provide accurate irradiance measurement data in a spectral range from 300 nm to 1100 nm (UV-Visible-NIR), with optical resolution FWHM < 7 nm, wavelength interval 0.3–0.5 nm wavelength accuracy ±0.2 nm, and field of view 180°. The EKO MB-22 is a prototype of RSB and has been commercially available since 2020 as the EKO RSB-01S. The rotating arm of the RSB is made of a 4 mm thick aluminum plate with a 25 mm band width and 285 mm radius, with a shielding angle of 5° full angle. In the first position (Irr. 1), the shadow band rests outside the field of view of the instrument; in the second position (Irr. 2), the shadow band stops at −5° from the sun disk; in the third position (Irr. 3), the RSB covers the sun disk to perform the DHI measurement; and finally, in the fourth position (Irr. 4), the shadow band stops at +5° from the sun disk. Thus, the irradiance components (GSI = DIR + DFR) were calculated. Observations were made at 5-minute intervals during the day. The whole-day GSI_(actual)_ was collected using the second-class pyranometer FMP3 (ISO9060:2018; Kipp & Zonen, The Netherlands) with a viewing angle of 180° in a wavelength range of 300–2800nm to confirm the relatability and consistency of the data. The glass domes of the radiometers were cleaned every week. To minimize the instrumental uncertainty, this study used the GSI as the quantum flux in a wavelength range from 200 to 1200 nm during the sunshine to sunset period measured using SBS to compute both the CWRs and the DF. In the shadow-band system used for this observation, there is a time delay in the observation due to band movement. Therefore, under conditions where the shadow effect of clouds varies greatly over time, such as on a lightly cloudy day, it may not be possible to obtain an appropriate DF. As the GSI_(actual)_, which measured the quantum fluxes from 300–2800 nm throughout the day and GSI_(300-1100)_ are highly correlated (R^2^*_adj_*= 0.97, Sup. 1g), the dataset was thoroughly inspected and suspicious data were removed before the analysis was performed.

The radiant energy *I* [W m^−2^ nm^−1^] was converted to photon flux *E*_QF_ [μmol m^−2^ s^−1^ nm^−1^] using the following formula (Eq. 6), which was developed according to Plank’s law:

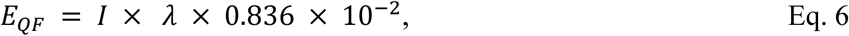

The photon flux in the critical wavelength range was integrated, and the CWRs were calculated for analysis. Data were analyzed using MATLAB to determine seasonal variations. The date was considered from a day, sunrise to sunset, and the percentage of dates was calculated as the percentage between the number of dates satisfying the given condition during the period and the total number of dates in the given period. To assess the statistical significance of the variation in DF and the observed CWRs across different seasons, we used the analysis of variance (*ANOVA*) and Tukey’s honestly significant difference post hoc tests to further elucidate our findings. The correlations between the variables were analyzed using Pearson’s correlation coefficient (*p*) and F-statistics (*F*) to gain a comprehensive understanding of variable interactions. Polynomial models were also generated to approximate the relationship between the DF and CWRs. The validity of these models was determined using the root mean square deviation (Eq. 7) and adjusted coefficient of determination (*R^2^_adj_*, Eq. 8):

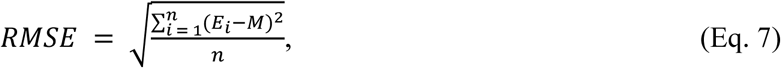

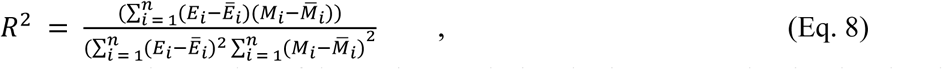

where *n* represents the number of data points, and *E*i and *M*i represent the simulated and measured values, respectively. The correlation of CWRs with AM and VP was tested, and the models were fitted using the optimum degree of polynomial fit. The model performance was evaluated similarly. Additionally, multiple linear models were fitted to test the combined effects of parameters on the CWRs.

## Results

### Diffuse fraction of solar radiation (DF)

The monthly mean DF varied between 0.58 and 0.79 (*std* > 0.2); the relative difference (*rd*) was 28.7% and the absolute difference (*ad*) was 35.27% (Table. 1). In addition, although no seasonal pattern in the daily mean DF was observed (Fig. 1a), there were significant differences in the daily mean DF by month (*F* = 2.24, *p* < 0.05). Specifically, in June, the rainy season, the daily mean DF was higher than 0.8, and in October, which is the autumn season, the daily mean DF was around 0.6. In addition, a wide range in daily mean DF was observed within each month throughout the year. When we categorized the DF values into five different groups by percentile range, we found that only approximately 10% of the annual data had a DF < 0.37, indicating clear-sky conditions. The majority (about 90%) of the dates in the year had a large DF, and more than 35% of the dates in the year had a very high daily mean DF (DF > 0.88) (Fig. 1b).

**Fig. 1.**
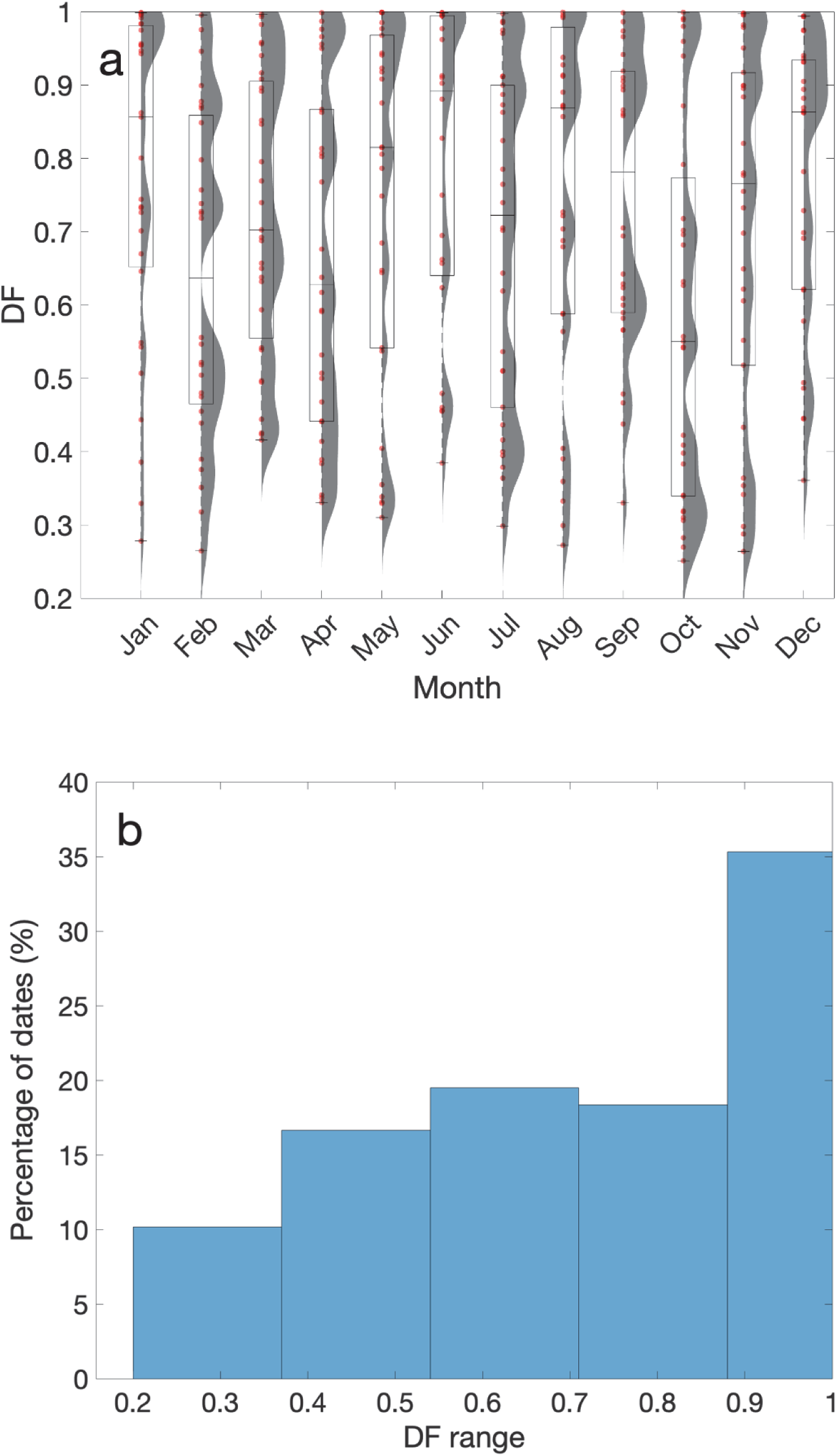
(**a**) Daily mean DF between months, raincloud plot with box plot constructed to observe the monthly variations, and box plot of daily mean DF **(b)** Relative frequency histogram of daily mean DF variation classifying the DF values into five different groups by percentile range to observe the distribution during 2021.

**Table. 1.** ANOVA statistics, absolute and relative differences CWRs, GSI, and DF between months.

| Wavelength ratio | Difference between the highest and lowest (%) |  | ANOVA results |
| --- | --- | --- | --- |
|  | Relative difference | Absolute difference | *F* statistics |
|  | (*rd*) | (*ad*) |  |
| GSI | 99.62 | 190.18 | 15.2*** |
| DF | 28.70 | 35.27 | 2.24** |
| R/B ratio | 5.74 | 5.91 | 6.36*** |
| R/G ratio | 2.99 | 3.04 | 9.26*** |
| B/G ratio | 2.55 | 2.59 | 4.59*** |
| R/PAR ratio | 2.85 | 2.89 | 6.88*** |
| B/PAR ratio | 2.72 | 2.76 | 5.68*** |
| G/PAR ratio | 0.36 | 0.36 | 14.59*** |
| R/FR ratio | 12.45 | 13.09 | 56.19*** |
| UVA/UVB ratio | 43.02 | 52.48 | 101.75*** |
| UV/PAR ratio | 10.90 | 11.52 | 7.31*** |
| UV/GSI ratio | 17.87 | 19.48 | 12.72*** |
| UV-B/B | 47.42 | 64.23 | 92.73*** |
| PAR/GSI ratio | 7.26 | 7.49 | 29.22*** |

\*\*\* $p < 0.001$ and \*\* $p < 0.01$

Seasonal relative frequency DF histograms were constructed using the same annual DF bin range to compare between the four seasons. The seasons were identified as winter (Dec, Jan, Feb), spring (Mar, Apr, May), summer (Jun, Jul, Aug), and autumn (Sep, Oct, Nov). The histograms show that the highest percentage of dates in each season fall into the DF > 0.88 category, which results in characteristically high seasonal daily mean DFs (Fig. 2). To test the similarity in the distribution of daily mean DF between seasons, we calculated the correlation of the percentage date distribution between the different DF ranges across seasons. A high correlation (*R^2^_adj_* = 0.76) was found between the winter and summer seasons concerning the percentage of days in each DF range, indicating a high seasonal similarity (Sup. 4). Conversely, autumn showed a distinctly different trend from all other seasons, with a much lower correlation (*R^2^_adj_* < 0.4) with the other seasons (Sup. 4). Winter and autumn showed the lowest correlation (*R^2^_adj_* = 0.1).

**Fig. 2.**
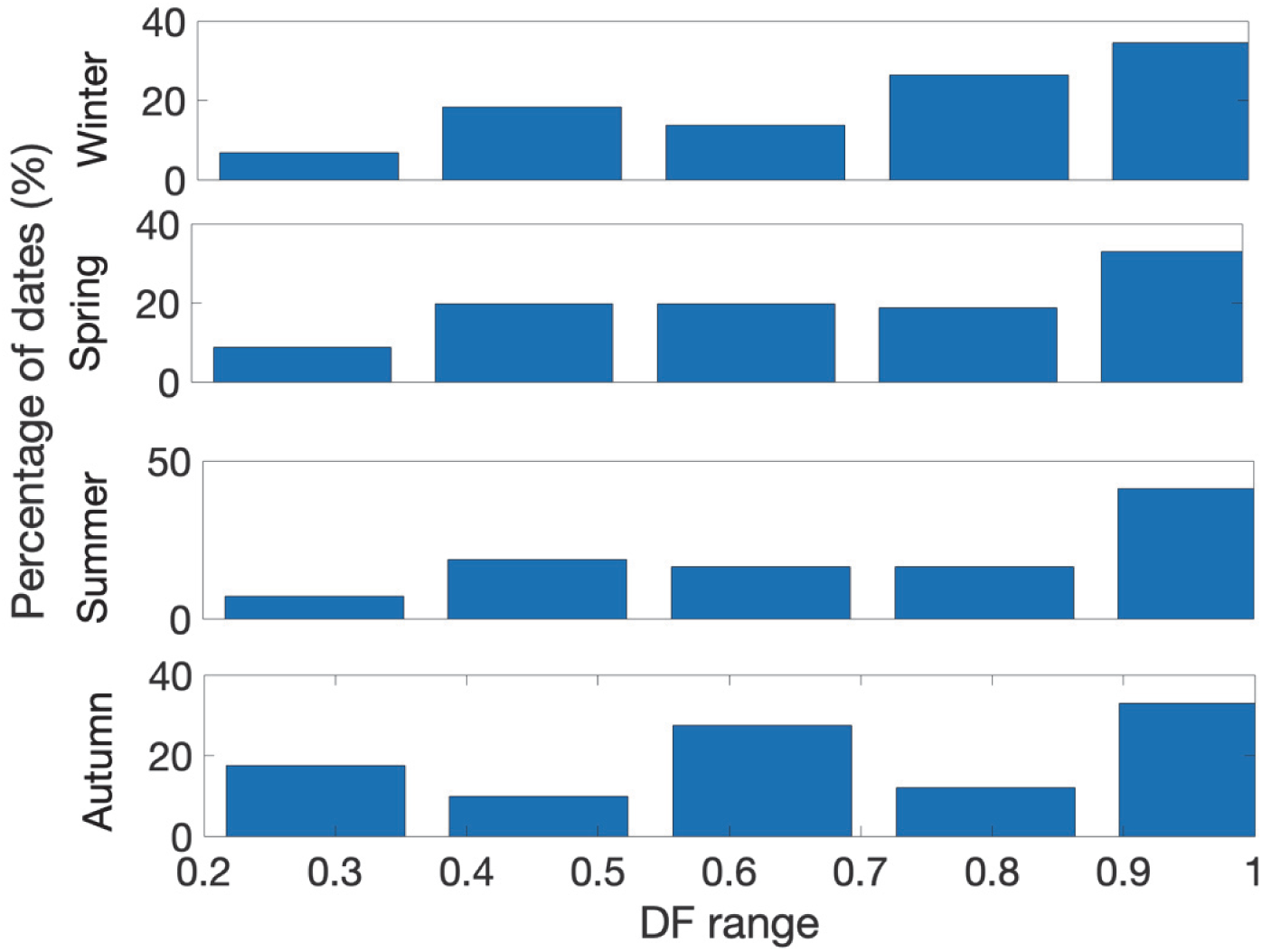
Seasonal subplot of the percentage of days within each DF range., The percentage of days distribution between different DF ranges among seasons was plotted for the same DF percentile range as Fig.1b, to observe the distribution similarity between different seasons during 2021.

### GSI and PAR

The monthly mean PAR/GSI_(300-1100)_ ratio varied significantly (0.43–0.47); it increased in summer and decreased in winter (*F* = 29.22, *p* < 0.001). The *rd* and *ad* of monthly average changes were 7.26% and 7.49%, respectively (Table. 1). The distribution pattern of PAR/GSI_(200-1200)_ showed a seasonal variation, characterized by an increase during the summer months and a decrease in the winter change in its distribution pattern, with an increase in summer and a decrease in winter (Fig. 3). The relationship between daily mean DF and daily mean GSI showed a significant negative correlation (*r* = −0.58, *p* < 0.001). GSI was approximated by a third-order polynomial with DF as a variable (*R^2^_adj_* = 0.34, Fig. 4).

**Fig. 3.**
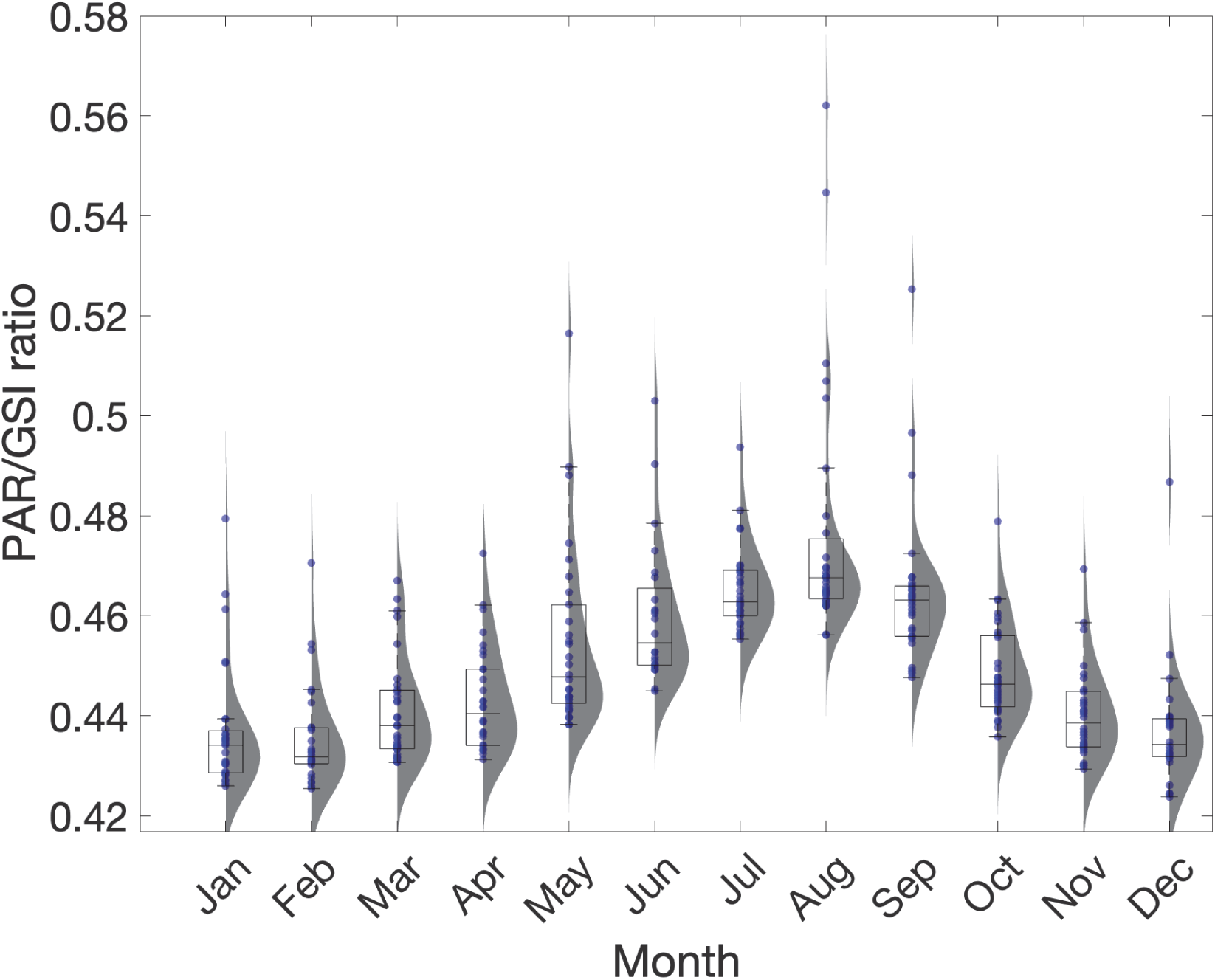
Daily mean PAR/GSI ratio by month, raincloud plot with box plot constructed to observe variations in daily mean DF by month during 2021.

**Fig. 4.**
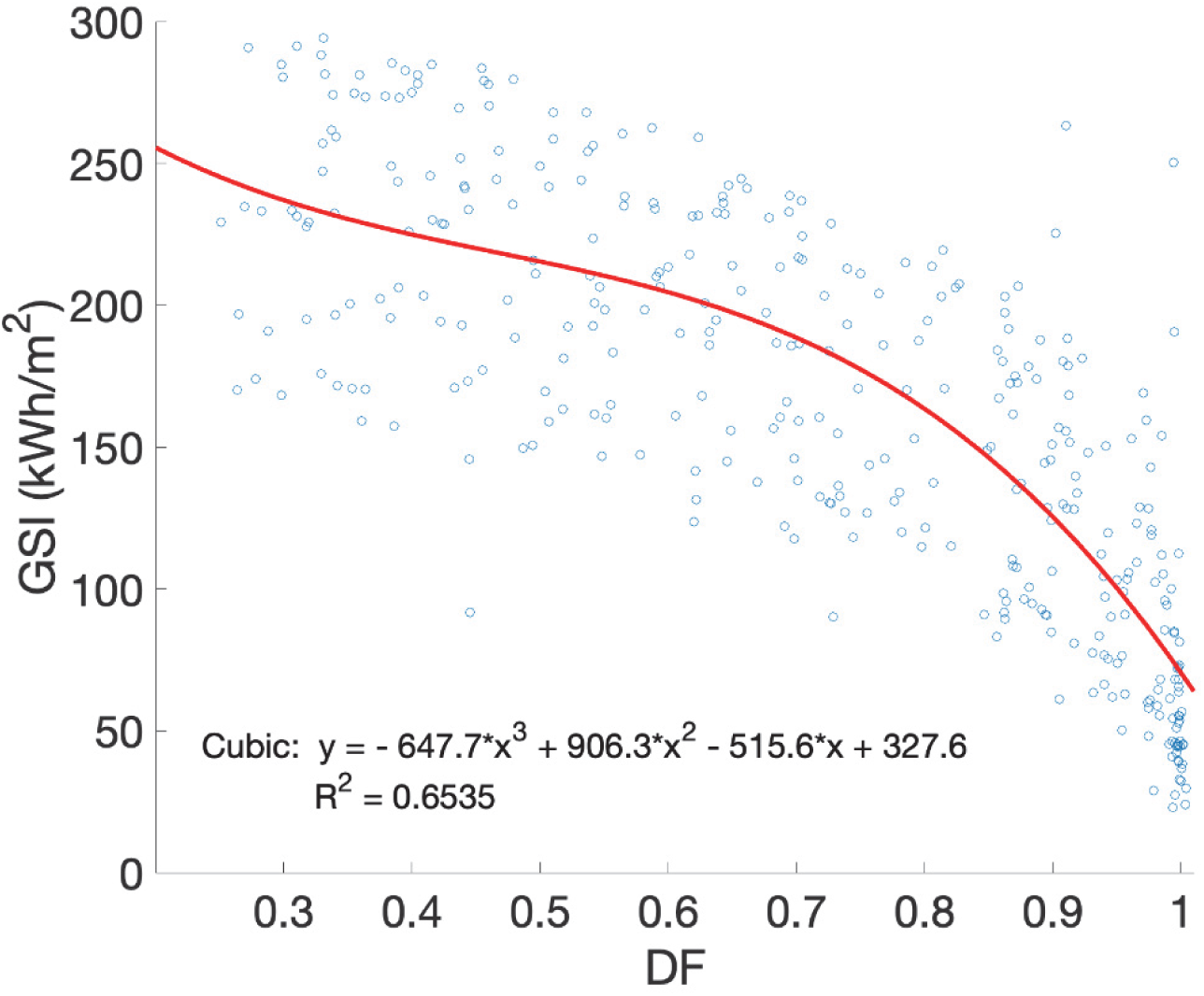
Daily mean GSI vs. daily mean DF, and the cubic model fitted to approximate the variation in GSI during 2021.

### PAR and UV

The seasonal variations in the UV ratios, especially the UV/PAR, UV-B/B, and UV-A/UV-B ratios, were analyzed. There are clear patterns in the annual variation of daily mean UV-B/B, UV/PAR (UV/GSI, Sup. 1f), and UV-A/UV-B ratios (Fig. 5).

**Fig. 5.**
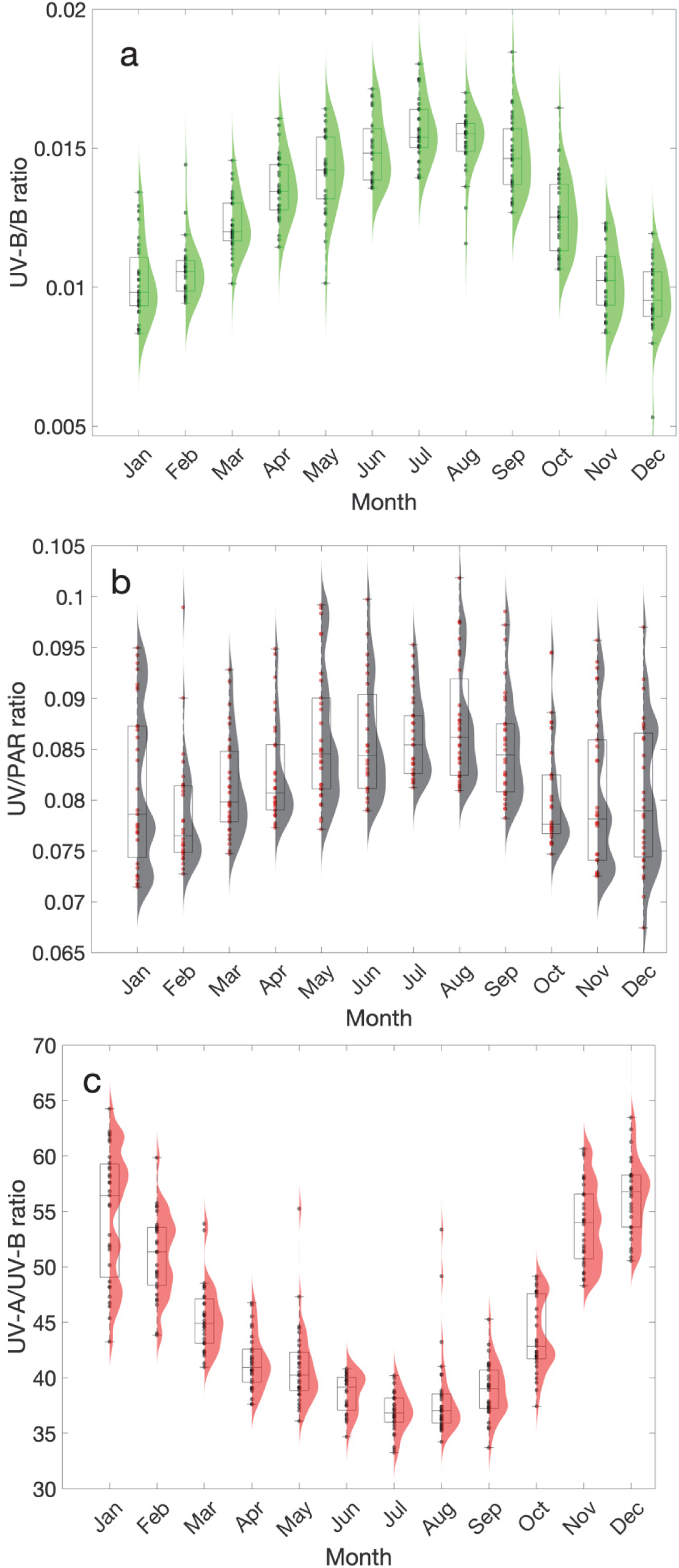
Annual variation of UV photon flux ratios, raincloud plot with box plot constructed to observe the variations in **(a)** daily mean UV-B/B ratios by month, **(b)** daily mean UV/PAR ratios by month, and **(c)** daily mean UV-A/UV-B ratios by month during 2021.

**Fig. 6.**
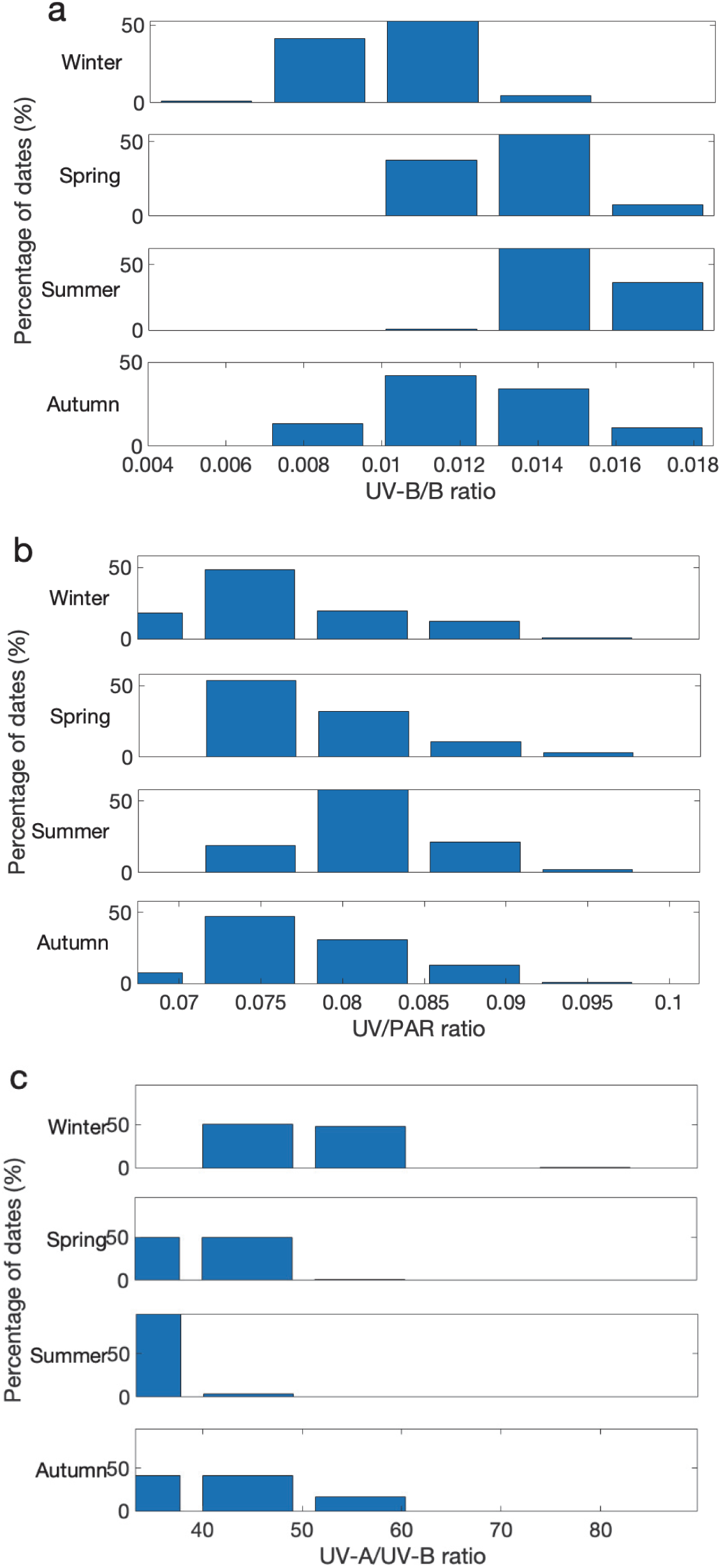
(**a**) Seasonal distribution of days within different ranges of UV-B/B ratio. **(b)** Seasonal distribution of days within different ranges of UV/PAR ratio. **(c)** Seasonal distribution of days within different ranges of UV-A/UV-B quantum flux ratio during 2021.

Significant differences were found in the daily mean UV-B/B ratio (*F* = 92.73, *p* < 0.001), UV/PAR ratio (*F* = 7.31, *p* < 0.001), and UV-A/UV-B ratio (*F* = 101.75, *p* < 0.001) between months. The *rd* of UV-B/GSI was 17.87%, and that of UV/PAR was 10.90%, which indicates a considerable variation (Table. 1). However, a very high *rd* was observed in the monthly mean of UV-A/UV-B and UV-B/B (43.02% and 47.42%, Table. 1). Furthermore, the UV-B/B and UV/PAR ratios behaved similarly (Fig. 5) by increasing in summer and decreasing in winter and autumn. In contrast, the UV-A/UV-B ratio behaved inversely by decreasing in summer and increasing in winter (Fig. 5a). The daily mean UV-B/B ratio showed seasonal uniqueness among the four seasons. Interestingly, summer stands out with a significant number of days in the highest UV-B/B ratio range (0.016–0.018), which is a feature not shared by other seasons. The significant correlation between spring and autumn (*R^2^_adj_* = 0.97) in percentage distribution of days between different UV/PAR ranges indicates a high degree of seasonal similarity in the UV/PAR ratio (Sup. 4). Furthermore, the significant correlation was found in the UV/PAR ratios between winter and autumn (*R^2^_adj_* = 0.83), suggesting similar behavior patterns. However, the percentage days distribution in summer was not correlated significantly with any of the other seasons, showing different characteristics, distinguishing it from the other seasons. UV/GSI ratio also behaves similarly (Sup. 4). Considering the percentage days distribution in the UV-A/UV-B ratio, only autumn and spring showed similarities (*R^2^_adj_* = 0.97) none of the other seasons had significant correlations in UV-A/UV-B ratios (*p* > 0.05, Sup. 4).

### PAR and RGB

PAR is of great importance in plant science. We analyzed the annual changes in wavelength ratios (CWRs) in the PAR wavelength range, especially R/B, R/G, and B/G ratios, which are important light signals for plant growth and development. The results showed a clear annual pattern of daily mean R/B and R/G ratios (Fig. 7a, b), which increased in winter and decreased in summer. In contrast, the daily mean B/G ratio increased in summer and decreased in winter (Fig. 7c).

**Fig. 7.**
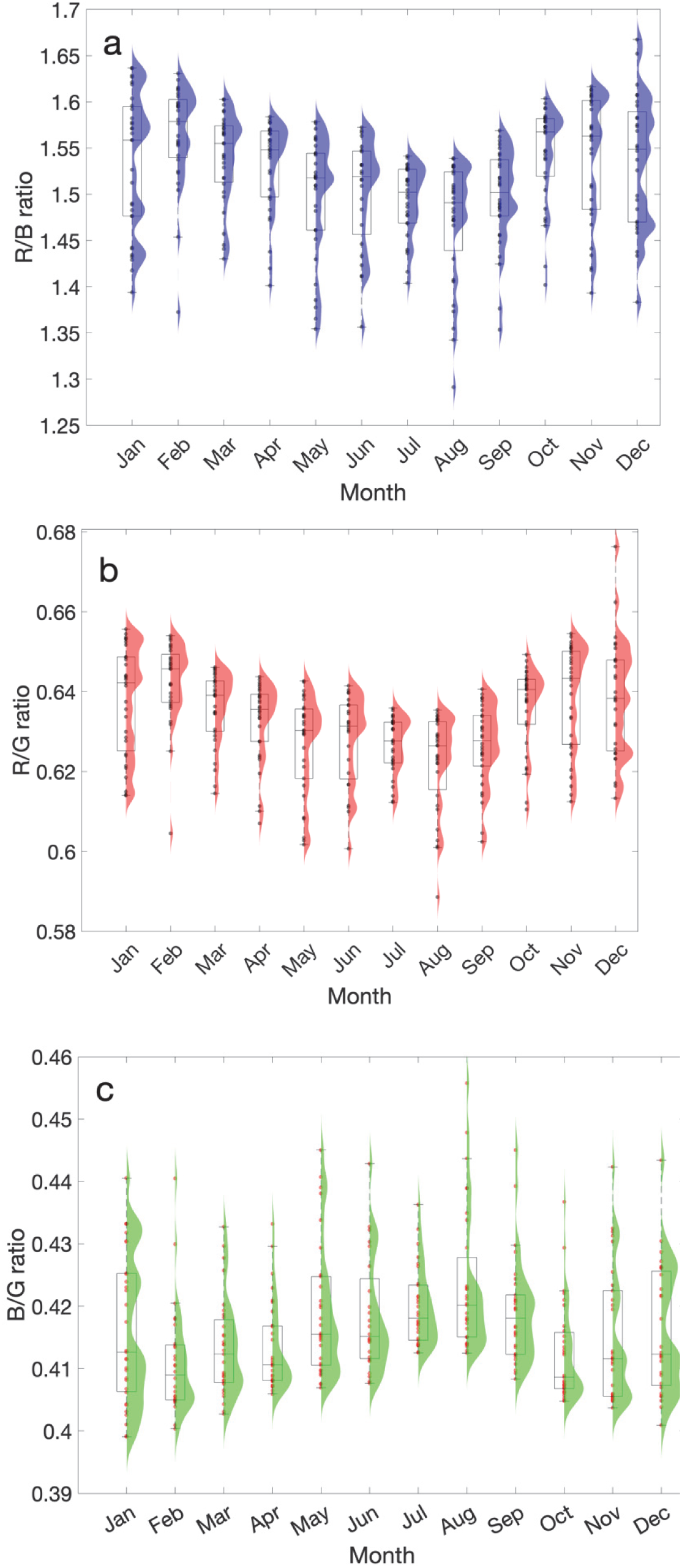
Annual fluctuation in R, B, and G wavelengths, raincloud plot with box plot constructed to observe the variations in (**a)** daily mean R/B quantum flux ratios by month, (**b)** daily mean R/G quantum flux ratios by month, (**c)** daily mean B/G quantum flux ratios by month during 2021.

There were significant seasonal changes in the R/B, R/G, and B/G ratios (Table. 1). However, the *rd*s of the monthly average R/B, R/G, and B/G ratios varied only slightly, at 5%, 2.7%, and 2.5%, respectively.

Winter, autumn, and spring were comparable in terms of the seasonal distribution of the daily average R/B ratio (Fig. 8a). The correlation between the R/B ratio distributions in autumn and spring was strong (*R^2^_adj_* = 1.00, *p* < 0.001, Fig. 8a, Sup. 4). The distribution of the daily mean R/G ratio was significantly different between winter and summer. Furthermore, the ratio in winter was comparable to that in autumn, and that in summer was comparable to that in spring (Fig. 8b). The seasonal daily mean B/G ratio showed no seasonal variation (Fig. 8c).

**Fig. 8.**
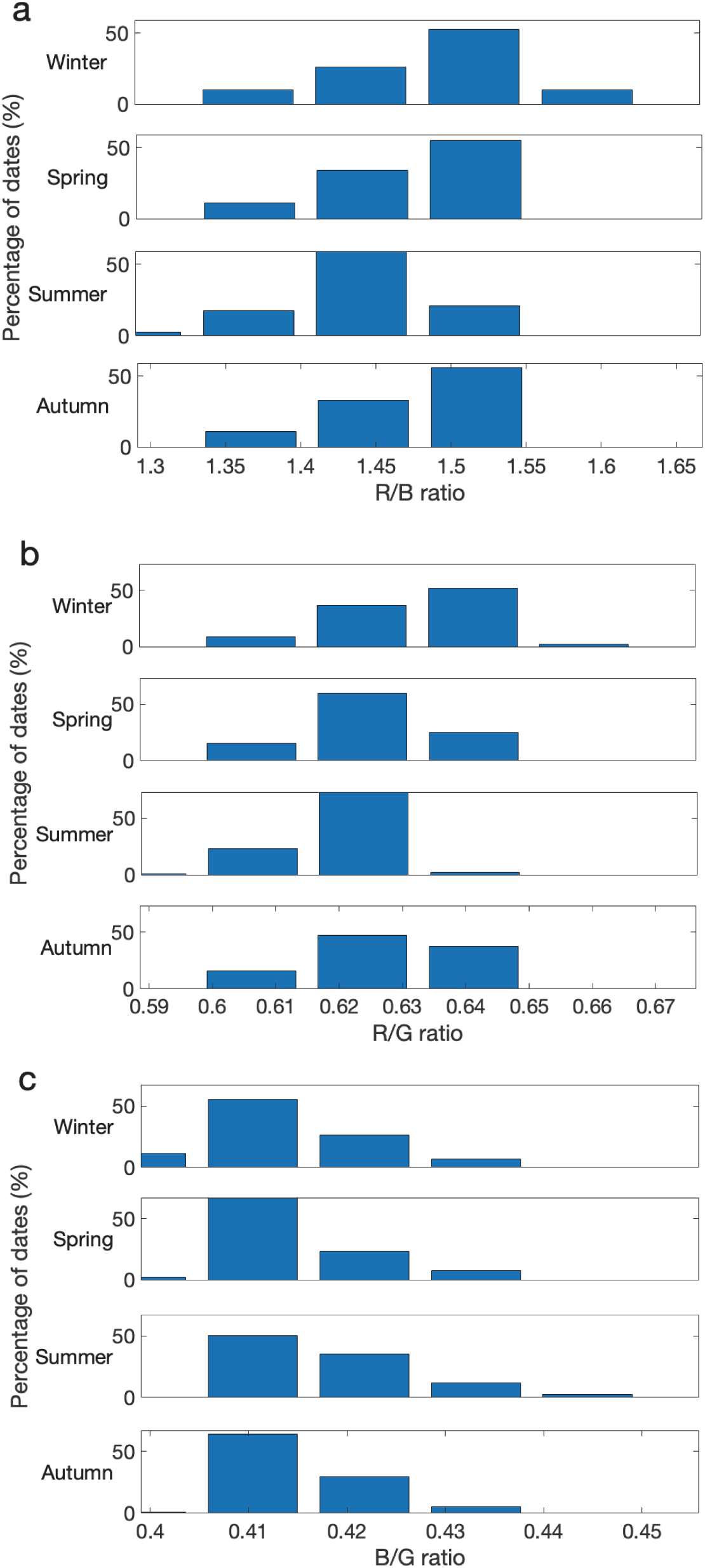
(**a**) Seasonal distribution of dates in different ranges of R/B photon flux ratio. **(b)** Seasonal distribution of dates in different ranges of R/G photon flux ratio. **(c)** Seasonal distribution of dates in different ranges of B/G photon flux ratio during 2021.

### R and FR

The R/FR ratio is a CWR recognized by plants and is widely discussed in plant physiological ecology. However, its annual and seasonal variations have received relatively little research attention. The daily average R/FR ratio increased in summer compared with that in winter (Fig. 9a), and significant monthly variation (*F* > 55, *p* < 0.001) was observed in the daily average R/FR ratio (Table. 1). Furthermore, the R/FR ratio showed significant changes of 12% *rd* and 13% *ads*. The R/FR ratio behaved differently in each season (Fig. 9b), and further interseasonal correlational analysis confirmed low correlation coefficients between seasons (*R^2^_adj_* < 0.50, *p* > 0.05, Sup. 4), except for fall and spring (*R^2^_adj_* = 0.84, *p* < 0.05, Sup. 4).

**Fig. 9.**
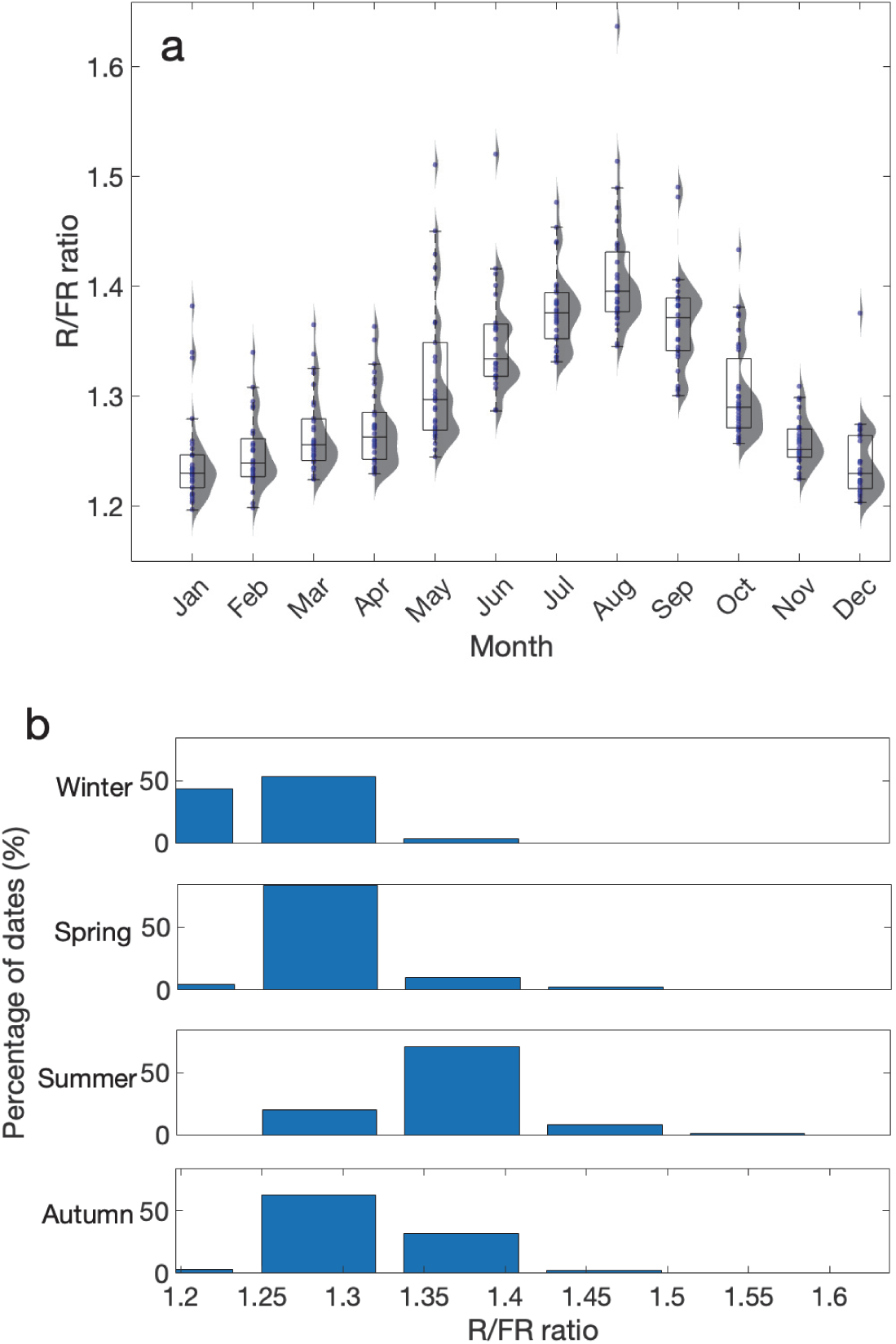
Annual fluctuation in R/FR photon flux ratio. (**a)** Daily mean R/FR quantum flux ratios by month, raincloud plot with box plot constructed to observe the variations, and **(b)** seasonal distribution of dates in different ranges of R/FR photon flux ratio during 2021.

### Effect of the atmospheric VP

The mean daily VP showed a strong positive correlation with the mean daily temperature (*r* = 0.94, *p* < 0.001, Table. 2). The constructed cubic model explained the variance between air temperature and VP (*R^2^_adj_* = 0.92, *RMSE* = 2.069, Fig. 10a). Furthermore, AM and VP showed a significant, strong, negative correlation (*r* = −0.75, *p* < 0.001, Table. 2), and the constructed linear model partially explained the relationship between the two variables (*R^2^_adj_* = 0.47, *RMSE* =5.8542, Fig. 10b, Table. 3). However, no significant correlation was found between DF and VP (*r* = 0.09, *p* = 0.092, Fig. 10c). The mean daily VP was significantly correlated with all of the CWRs considered (*p* < 0.001, Table. 2). However, the strength of all the correlations was weak (|r| < 0.5, Table. 2), except for the UV-A/UV-B, R/FR, PAR/GSI, and UV-B/B ratio.

**Table. 2.** Person correlation statistics for the relationship between daily mean CWRs and daily mean DF and AM.

| Parameters | CWR/ Parameter | Pearson correlation coefficient |  |  |
| --- | --- | --- | --- | --- |
|  |  | DF | AM | VP |
| CWR | UV/PAR ratio | 0.76*** | −0.35*** | 0.44*** |
|  | UV/GSI ratio | 0.67*** | −0.42*** | 0.58*** |
|  | UV-A/UV-B ratio | 0.04 | 0.88*** | −0.76*** |
|  | PAR/GSI | 0.40*** | −0.51*** | 0.76*** |
|  | R/B ratio | −0.76*** | 0.29*** | −0.42*** |
|  | R/G ratio | −0.71*** | 0.41*** | −0.48*** |
|  | B/G ratio | 0.76*** | −0.17** | −0.48*** |
|  | R/FR ratio | 0.30*** | −0.59*** | 0.88*** |
|  | UV-B/B | 0.26*** | −0.82*** | 0.81*** |
| Other Parameters | DF |  | −0.003 | 0.09 |
|  | AM |  |  | −0.69*** |
|  | VP |  |  |  |
|  | Temperature | −0.05 | −0.76*** | 0.94*** |

\*\*\* $p < 0.001$ and \*\* $p < 0.01$

**Fig. 10.**
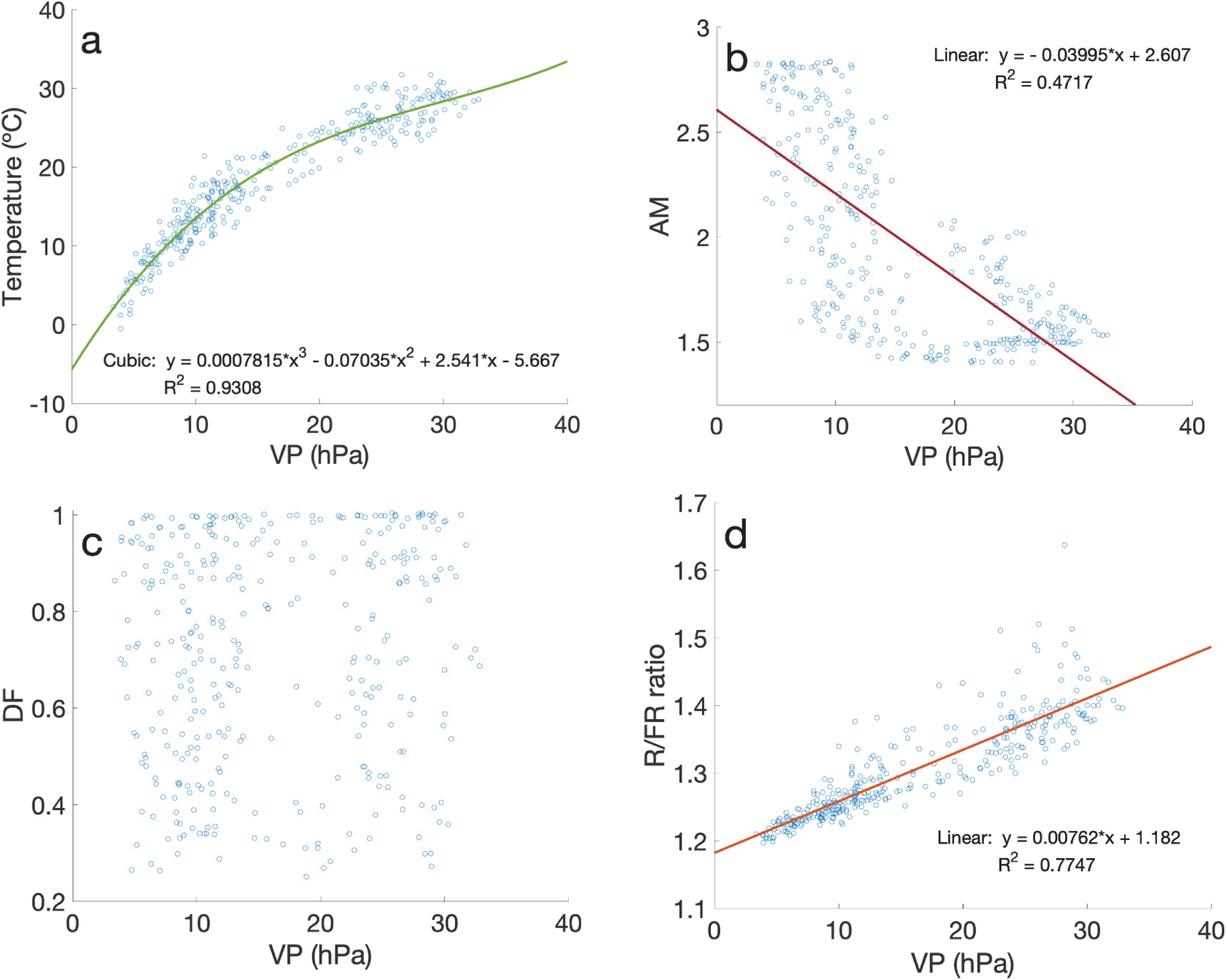
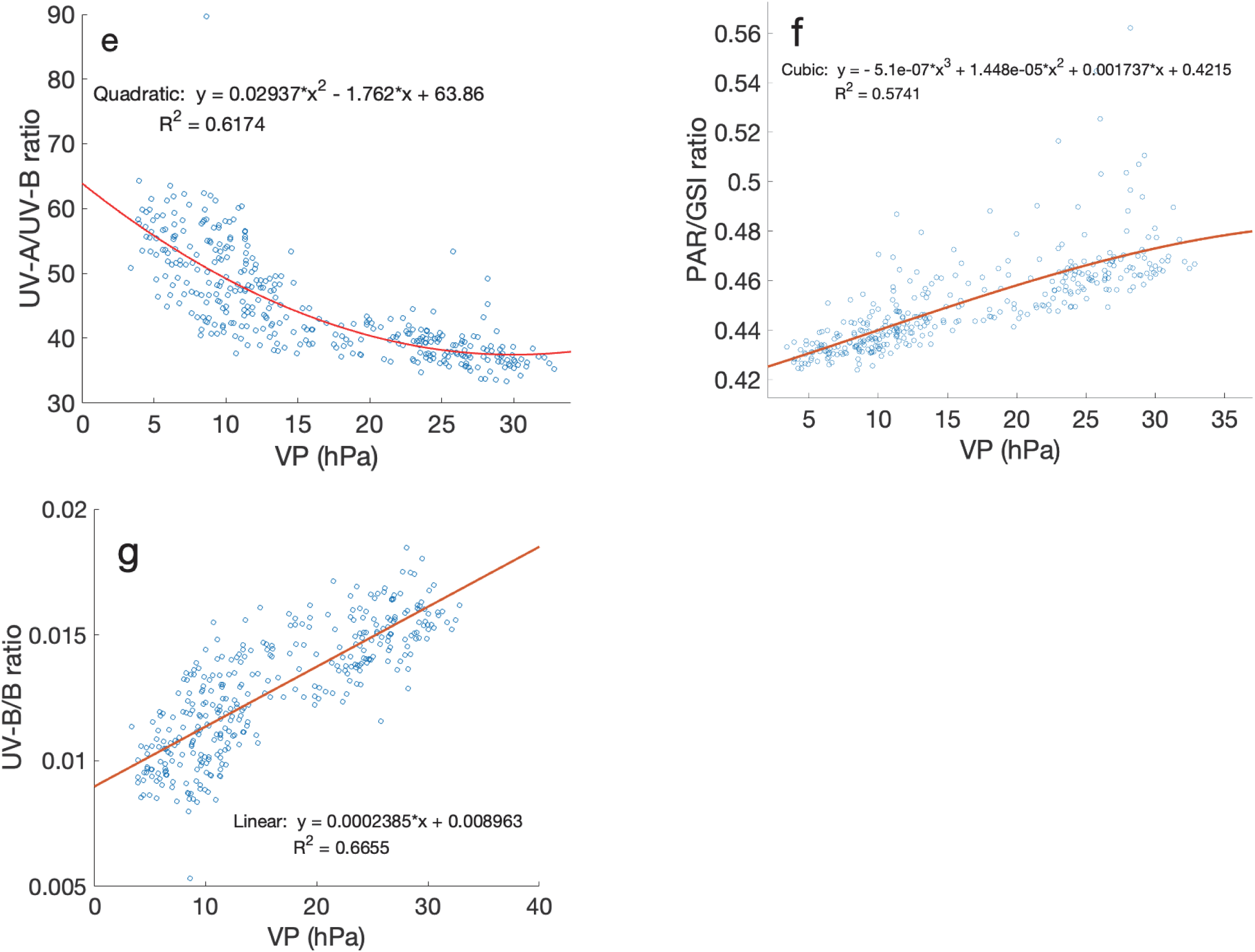
(**a**) Daily mean temperature vs. daily mean VP., and the cubic model constructed to approximate the VP. **(b)** Daily mean VP vs. daily mean AM, and the linear model constructed to approximate **(c)** daily mean DF vs. daily mean VP, **(d)** daily mean VP vs. daily mean R/FR photon flux ratio. The linear model constructed to approximate **(e)** daily mean VP vs. daily mean UV-A/UV-B photon flux ratio, and quadratic model constructed to approximate **(f)** daily mean VP vs. daily mean PAR/GSI photon flux ratio. The linear model constructed to approximate **(g)** daily mean VP vs. daily mean UV-B/B photon flux ratio, and the linear model constructed to approximate during 2021.

The VP correlated positively with the UV-B/B, PAR/GSI and R/FR ratios (*r* > 0.75, *p* < 0.001). Conversely, it had a strong negative correlation with the UV-A/UV-B ratio (*r* = −0.76, *p* < 0.001, Table. 2). The constructed linear model for the R/FR ratio using VP explained 77% of its variation (*R^2^_adj_* = 0.77, *RMSE* = 0.034, Fig. 10d, Table. 3), and the constructed quadratic model for the UV-A/UV-B ratio using VP explained 61% of its variation (*R^2^_adj_* = 0.6152, *RMSE* = 4.94, Fig. 10e, Table. 3). Furthermore, the cubic model constructed for the PAR/GSI ratio using the VP explained 57% of its variation (*R^2^_adj_* = 0.57, *RMSE* = 0.0123, Fig. 10f, Table. 3), and the constructed linear model for the UV-B/B ratio using VP explained approximately 65 % of its variation (*R^2^_adj_* = 0.67, *RMSE* = 0.00023, Fig. 10g, Table. 3). However, the VP alone could not explain the variation in the other CWRs (*R^2^_adj_* < 0.25).

**Table. 3.**
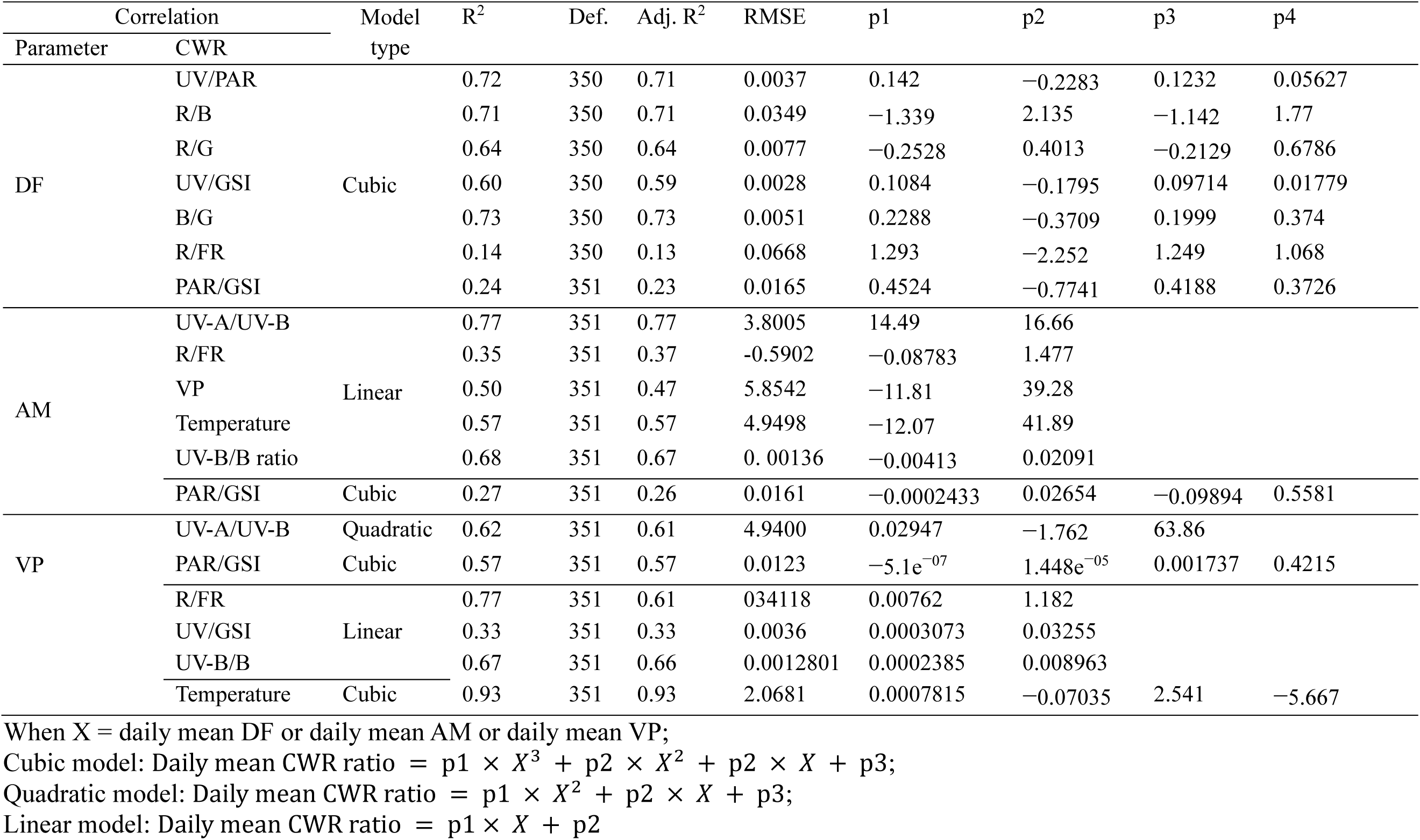
The model summaries of the polynomial models for daily mean CWRs vs. daily mean DF, daily mean AM, and daily mean VP.

### Effect of AM on CWRs

The daily mean AM showed a strong significant negative correlation with temperature (*r* = −0.75, *p* < 0.001, Table. 2), and a correlation with VP. The constructed linear model partly explained the variance between air temperature and AM (*R^2^_adj_* = 0.56, *RMSE* = 4.9498, Table. 3, Fig. 11a). However, no significant correlation was found between AM and DF (*r* < 0.006, *p* > 0.05, Table. 2). All the considered CWRs were significantly correlated with the daily mean AM (*p* < 0.001, Table. 2). However, the correlations were weak (|r|< 0.3, Table. 2), except for the UV-A/UV-B, R/FR, PAR/GSI, and UV-B/B ratios. The daily mean AM showed a strong correlation with the daily mean UV-A/UV-B ratio (*r* = 0. 88, *p* < 0.001, Table. 2) and the linear model developed using AM was able to explain more than 75% of the variation in the daily mean UV-A/UV-B ratio (*R^2^_adj_* = 0.77, *RMSE* = 3.800, Table. 3, Fig. 11b). The AM was negatively correlated with the R/FR and PAR/GSI ratios (*r* < −0.50, *p* < 0.001, Table. 2). The AM did not well explain the variation in the R/FR and PAR/GSI ratios in linear or cubic models (Table. 3, Fig. 11c, d). The daily mean AM showed a strong negative correlation with the daily mean UV-B/B ratio (*r* = −0. 82, *p* < 0.001, Table. 2), and the linear model developed using AM was able to explain over 67% of the variation in the daily mean UV-B/B ratio (*R^2^* = 0.77, *RMSE* = 0.001, Table. 3, Fig. 11e).

**Fig. 11.**
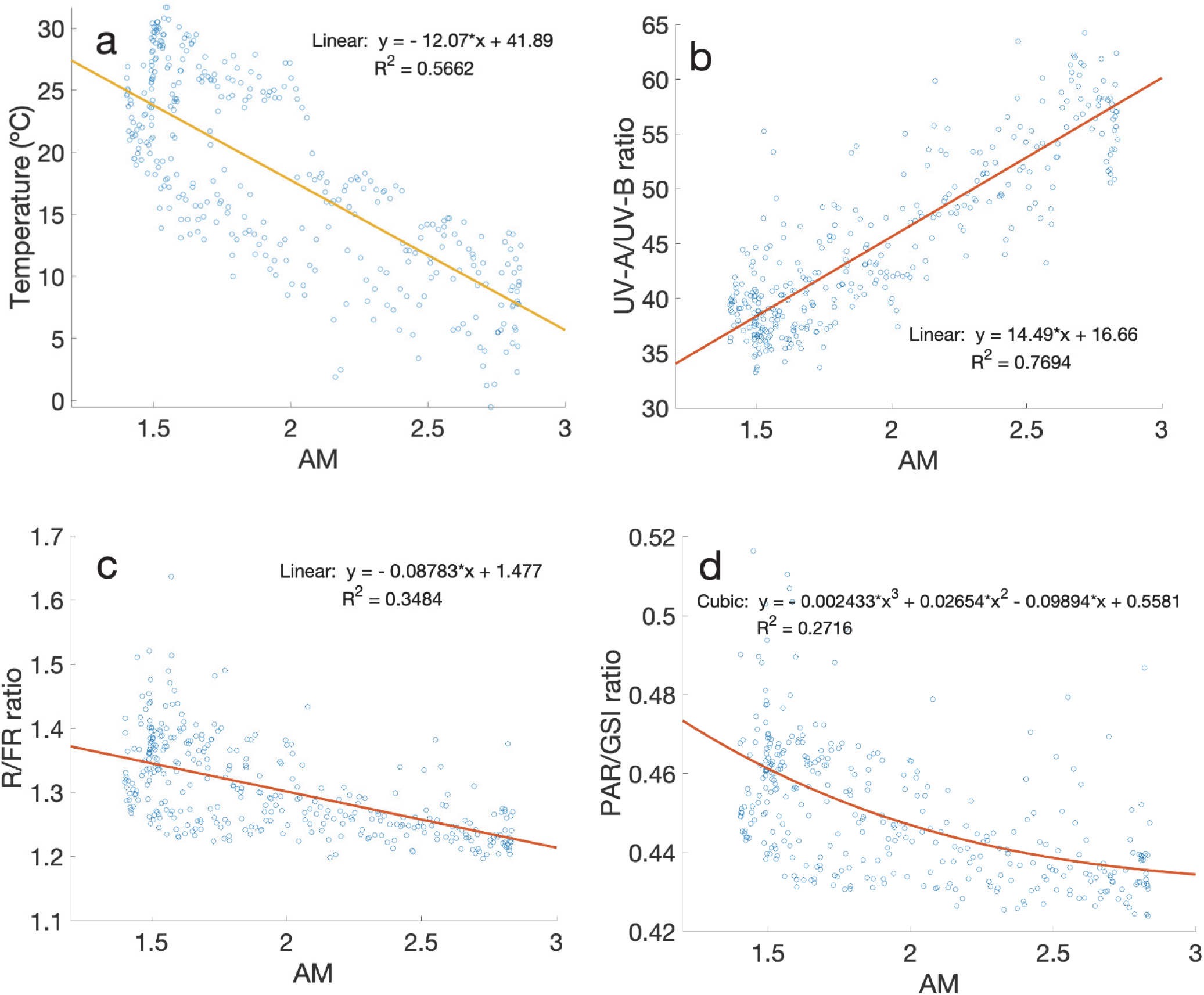
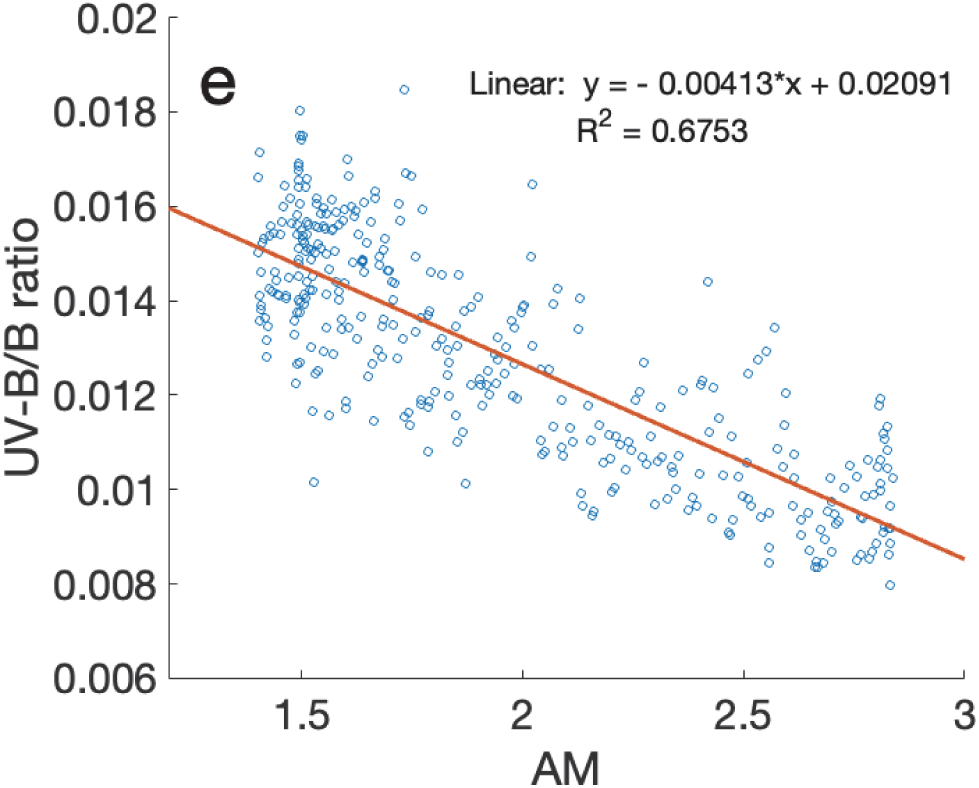
(**a**) Daily mean temperature vs. daily mean AM, and the linear model constructed to approximate **(b)** daily mean AM vs. daily mean UV-A/UV-B photon flux ratio; daily mean VP vs daily mean AM, and the linear model constructed to approximate **(c)** daily mean AM vs. daily mean R/FR photon flux ratio. The linear model constructed to approximate **(d)** daily mean AM vs. daily mean PAR/GSI photon flux ratio, and the cubic model constructed to approximate **(e)** daily mean AM vs. daily mean UV-B/B photon flux ratio, and the linear model constructed to approximate during 2021.

### Effect of DF on CWRs

The daily mean DF did not show a significant correlation with UV-A/UV-B (*p* > 0.05, Table 2, Fig. 12a). However, DF was strongly correlated with most of the CWRs, including UV/PAR, R/B, R/G, and B/G (Table 2). Additionally, the UV/PAR and B/G ratios were positively correlated with DF (*r* > 0.70, *p* < 0.001), whereas the R/B and R/G ratios were negatively correlated with DF (*r* < −0.70, *p* < 0.001). The UV/GSI ratio had a positive and significant correlation with DF (*r* = 0.67, *p* < 0.001), whereas the R/FR and UV-B/B ratio had a weak but significant correlation with DF (*r* = 0.30, *p* < 0.001, Fig. 12). A cubic polynomial model can be used to approximate the relationship between UV/PAR, R/B, B/G, and daily mean DF (*R^2^_adj_* > 0.70, *RMSE* < 0.0349, Table. 3). Cubic polynomial models with DF as a variable were used to accurately approximate the daily mean ratios of R/G and UV/GSI (*R^2^_adj_* > 0.59, *RMSE* < 0.0077, Table. 3). The daily mean ratios of UV/PAR, R/B, R/G, B/G, and UV/GSI were modeled using the daily mean DF as a variable (Table. 3, Fig. 12).

**Fig. 12.**
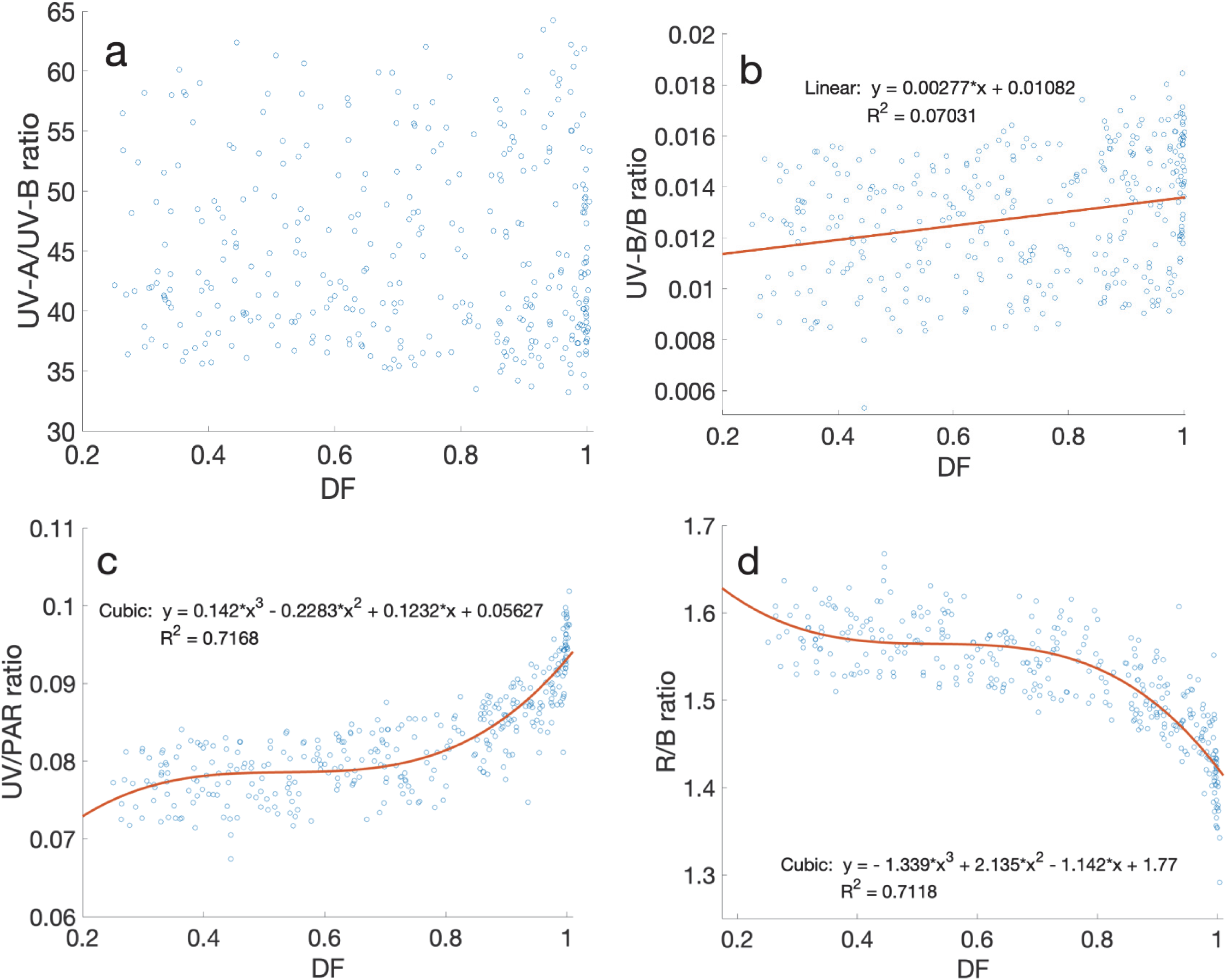
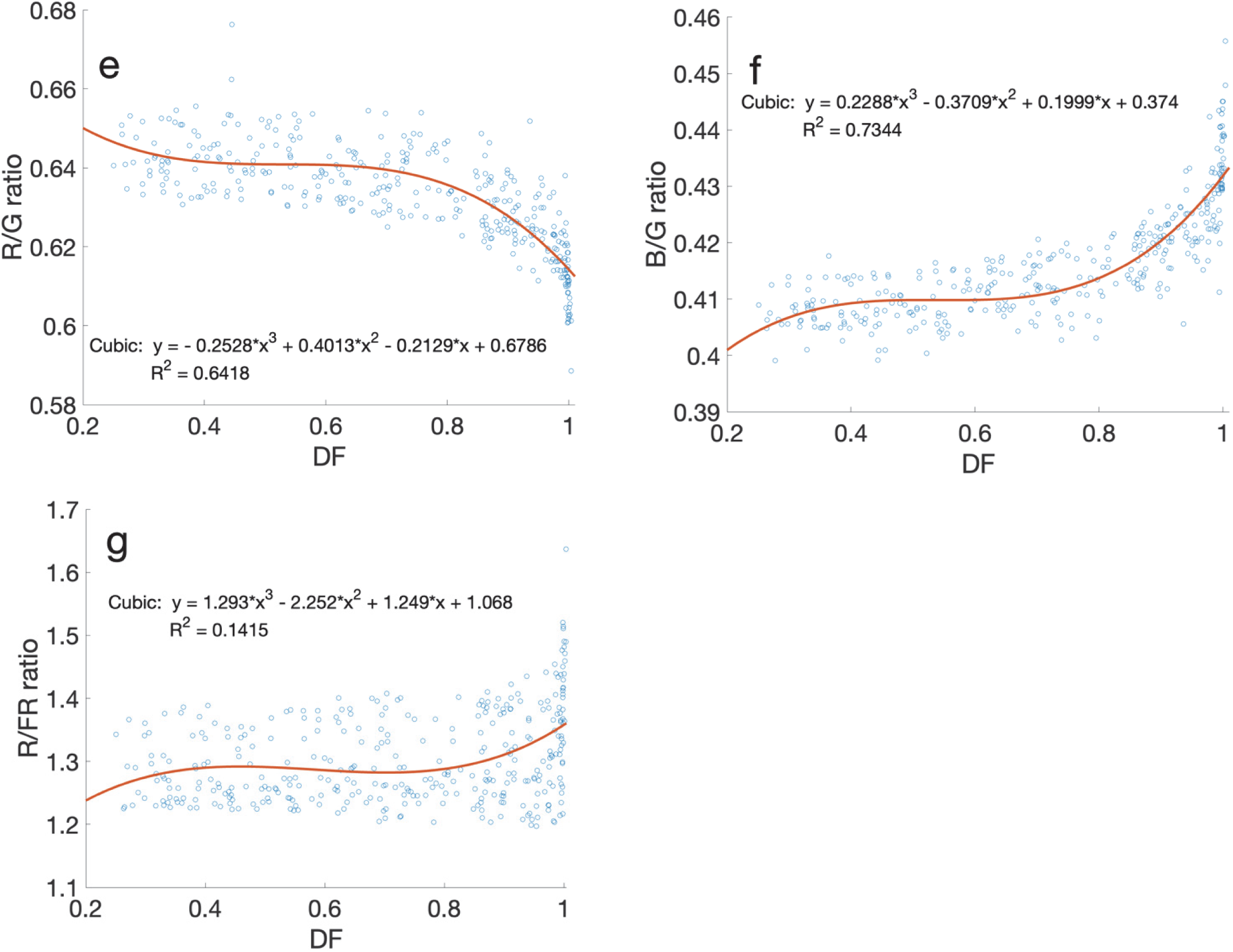
Relations between daily mean DF vs. CWRs and the cubic model constructed to approximate the variations in **(a)** daily mean UV-A/UV-B quantum flux ratio vs. daily mean AM, **(b)** daily mean UV-B/B quantum flux ratio vs. daily mean DF, **(c)** daily mean UV/PAR quantum flux ratio vs. daily mean DF, **(d)** daily mean UV/GSI quantum flux ratio vs. daily mean DF, **(e)** daily mean R/B quantum flux ratio vs. daily mean DF. **(f)** daily mean R/G quantum flux ratio vs. daily mean DF. **(g)** daily mean B/G quantum flux ratio vs. daily mean DF. **(h)** daily mean R/FR quantum flux ratio vs daily mean DF during 2021.

### Combined effects of parameters

To evaluate the effects of multiple variables, approximate models were created by multiple linear regression. Models were constructed using the combination of considered three parameters (VP, AM, and DF) to compute the CWRs, and the best ones were selected. The VP and AM together explained 82% of the total variation in the UV-A/UV-B ratio (*R^2^_adj_* = 0. 8231, *RMSE* = 3.8005, Table. 4, Fig. 13a). The VP and DF together explained more than 80% of the variation in the R/FR ratio (*R^2^_adj_* = 0.82, *RMSE* = 0.0311, Table. 4, Fig. 13b). The VP and DF together explained approximately 69% of the total variation in the PAR/GSI quantum flux ratio (*R^2^_adj_* = 0.69, *RMSE* = 0. 0101, Table. 4, Fig. 13c). Moreover, the AM, VP and DF together explained approximately 84% of the variation in UV-B/B ratio (*R^2^_adj_* = 0.84, *RMSE* = 0. 0009, Table. 4, Fig. 13d).

**Fig. 13.**
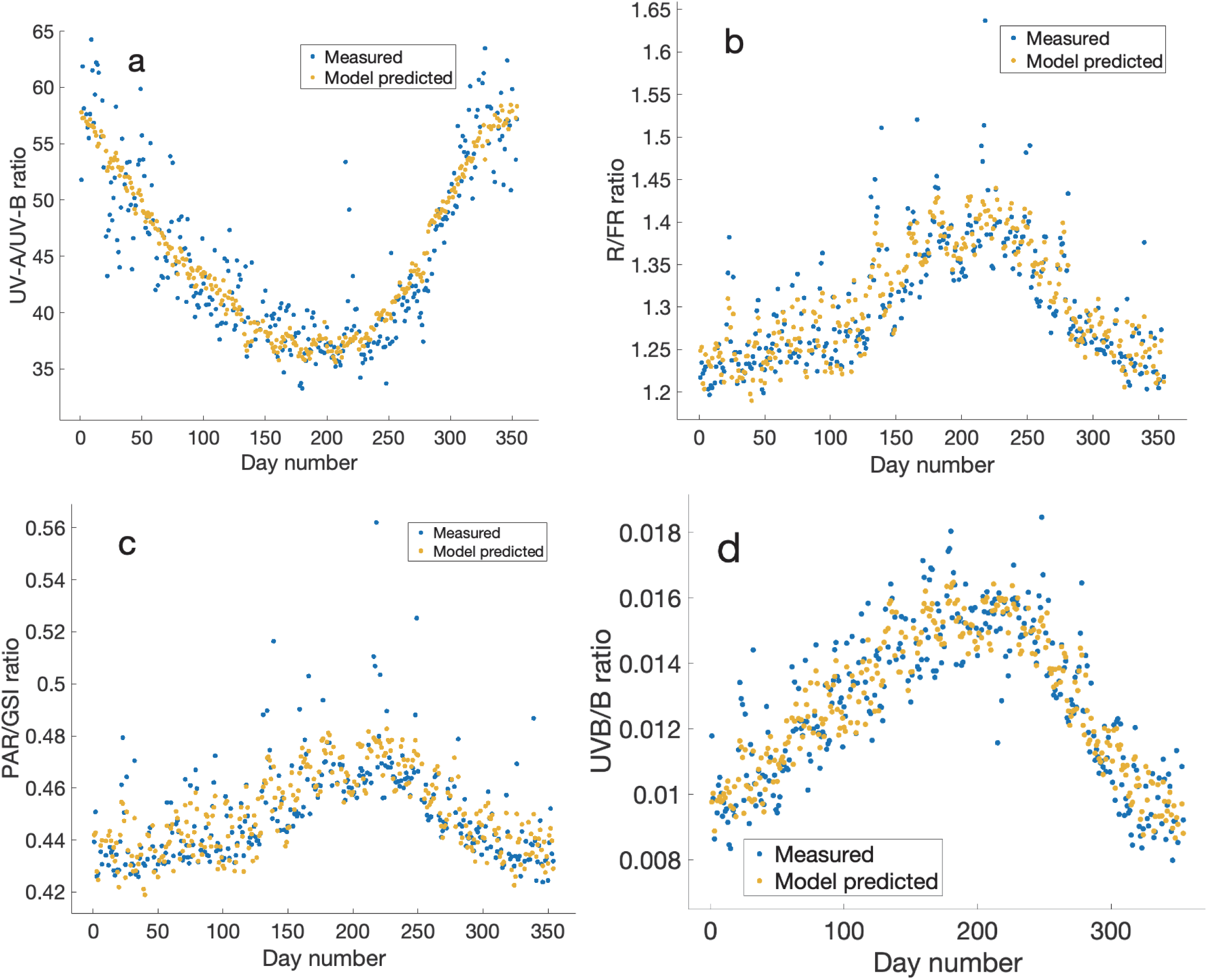
Comparison of measured and predicted multiple linear model outputs for the measured period. The x-axis indicates the day of the year in 2021. **(a)** Model developed using VP and AM to predict the UV-A/UV-B ratio on the y-axis. **(b)** Model developed using VP and DF to predict the R/FR ratio on the y-axis. **(c)** Model developed using VP and DF to predict the PAR/GSI ratio on the y-axis. **(d)** Model developed using DF, AM and VP to predict the UV-B/B ratio on the y-axis during the year 2021.

**Table. 4.**
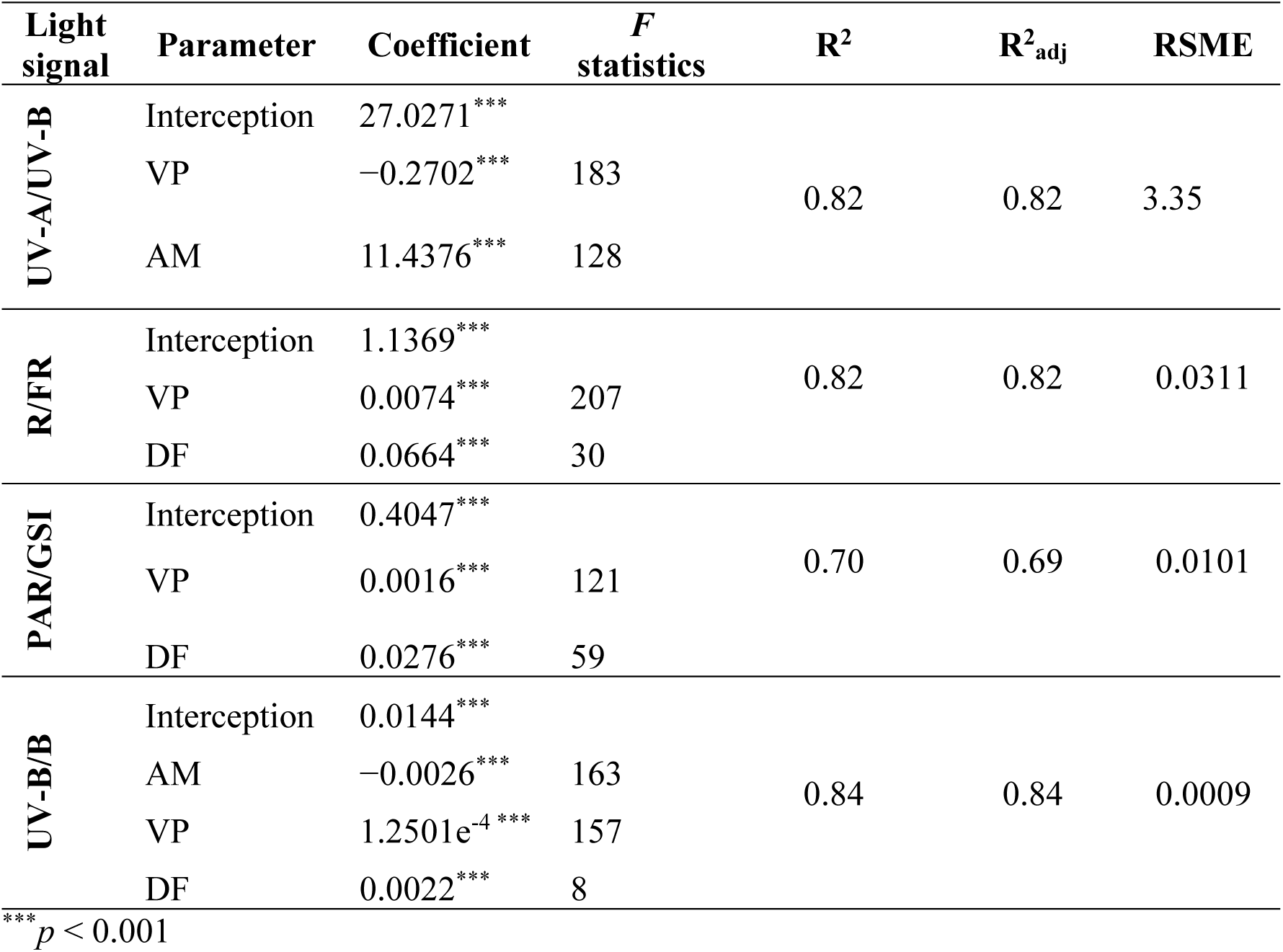
Multiple linear model summaries of the combined effects of the parameters VP, AM, and DF on specific CWRs.

## Discussion

In the midlatitude zone, which includes East Asia, strong westerly winds (jet streams) in the upper atmosphere move north and south and are influenced by the tropics in summer and by the cold zone in winter; therefore, the difference between the seasons is quite clear. In addition, East Asia, including Japan, is located to the east of the Eurasian continent and has a monsoon climate, which is strongly influenced by seasonal winds (monsoons) blowing near the ground due to the difference in specific heat between the continent and the ocean. The climate where the Itoshima Peninsula is located, is characterized by strong seasonal differences, with warm, humid seasonal winds blowing from the south in summer and cold seasonal winds blowing from the north in winter (Yoshino and Kazuko 1981). These seasonal differences significantly affect the local atmospheric environment and change the nature of the incident SR.

Generally, PAR decreases as DF increases with increasing cloud cover. However, Liu et al. (2022) found that total primary production (GPP) increased as DF increased to 0.6, despite the decrease in PAR. Additionally, light use efficiency (LUE) significantly increased with an increasing DF (Liu et al. 2022). Because our results showed high DF in all seasons (Fig. 1a), using a clear sky model to compare plant responses may significantly underestimate GPP and LUE. Furthermore, evapotranspiration (ET) increases more than does direct light with an increasing DF for most plants (Wang et al. 2022). Because most days on the Itoshima Peninsula had a high DF (Fig. 1b), it is expected that ET may increase and NEP may be higher than on sunny days for most of the year. The average monthly DF varied widely, with a monthly *ad* of 35%. This variability may affect plant NEP, because diffuse light reduces foliar photosynthesis by 10% – 15%, but promotes NEP under low light conditions (Brodersen et al. 2008). Furthermore, most plants that were exposed to DF > 0.8 exhibited a 1.5-fold increase in ET unlike those exposed to direct light (DF < 0.2) when PAR < 300 Wm^−2^ (Wang et al. 2022). Our results highlight the limitations of the clear-sky model in relation to plant ecophysiology.

Contrary to previous assumptions that winter and summer are significantly different with respect to DF (Li and Yang 2015), our results showed that winter and summer DF conditions are not considerably different in terms of the percentage of days distributed in each DF range in each season (Fig. 2). However, we observed large variations (DF > 0.7 range) in the daily mean DF for most months (Fig. 1a), which indicated that SR diffusion varies significantly with the daily weather conditions. A fluctuating light intensity may lead to lower photosynthetic efficiency (Shi et al. 2022) and preferentially damage PS I compared with a consistently intense light (Kono et al. 2014; Shi et al. 2022). Large fluctuations in DF that lead to changes in light intensity could trigger a defensive photosynthetic response for most days and months. Because of the moderately strong correlation between daily mean DF and daily mean GSI (*r* = −0.58, *p* < 0.001; Fig. 4), both factors can work together to bring about significant physiological changes in plants. Therefore, it is crucial to consider both factors from a plant ecophysiological perspective. Field studies involving photosynthesis often multiply GSI by a constant (0.45 to 0.46) to estimate PAR. However, our results showed significant variation (*p* < 0.001, *F* = 29.22) in the monthly mean PAR/GSI_(300-1200)_ ratio (0.43 to 0.47; Fig. 3), which is consistent with the findings of a previous study (Akitsu et al. 2022).

UV radiation is mainly absorbed by the stratospheric ozone layer in the atmosphere, which is strongly influenced by the AM. Plant light signaling is influenced by UV/PAR ratios (Rai et al. 2021) and UV-A/UV-B ratios (Neugart and Schreiner 2018). Changes in UV-related CWRs may affect plant growth and development by altering the light signals of plants (Kong and Okajima 2016; Rai et al. 2021b; Jadidi et al. 2023). Our results showed clear seasonal variations in the UV/PAR, UV/GSI, and UV-A/UV-B ratios (Fig. 5). The UV/PAR and UV/GSI ratios increased during summer and decreased significantly during winter (Fig. 5a, Sup. 1f). The UV-A/UV-B ratios decreased significantly during the summer (Fig. 5c). Understanding the seasonal variation in UV-related CWRs it is crucial because the variations in UV-related ratios affect numerous plant functions (Yadav et al. 2020). As shown by Tissot and Ulm (2020), UV-B and B are sensed by cross-regulated photoreceptors in a polychromatic light environment. We have elucidated that VP and DF contribute to the enhancement of the UV-B/B ratio, while AM is associated with its attenuation.

Wavelengths in the PAR range of SR are a source of energy for photosynthesis and play an important role in light signal transduction in plants. The R/B ratio (Wang et al. 2022), the R/G ratio (Trojak et al. 2022), and the B/G ratio (Sellaro et al. 2010b) in addition to the R/FR ratio (Tan et al. 2022) have been reported as wavelength ratios that are effective for plant light perception. Although the variation in CWRs such as the R/B ratio (*ad* and *rd* ≈ 5.7%), R/G ratio (*ad* and *rd* ≈ 3%), and B/G ratio (*ad* and *rd* ≈ 2.5%) were not as large as other CWRs, these changes were statistically different (Table. 1, Sup. 3.). Our results showed that the R/B ratio varied between 1.3 and 1.7 (Fig. 7a). Since the plant are highly sensitive to light signals this change could effect on plant and ecosystem Considering the results of previous studies, an increase in the R/B ratio within the range of 1 to 2 may lead to an increase in fresh weight, chlorophyll content, water use efficiency, and land use efficiency in winter (Pennisi, Blasioli, et al. 2019; Pennisi, Orsini, et al. 2019). Because plants are highly sensitive to light signals, these changes in CWRs may be perceived as light signals by plants.

The R/G ratio decreased in summer and increased in winter (Fig. 7b). Although there have been inconsistent reports on the effect of this photon ratio, the ratio may serve as a signal that influences plant height and leaf characteristics (Park and Runkle 2023). In the field environment, there is a considerable increase in the summer R/FR ratio (*ad* ≈ 13%); therefor, there may be a combined effect with the summer R/FR ratio and the summer R/G ratio (Fig. 9). The B/G ratio has been reported to affect plant shade response, (Liu and van Iersel 2021). However, although the change in the B/G ratio in the field was statistically significant (*p* < 0.001), the variation was minimal (*ad* and *rd* ≈ 2.5%; Fig 7c). Therefore, seasonal effects on the B/G ratio may not occur. Instead, plants may use the B/G ratio as a signal to detect shade from the canopy.

It has been reported that the R/FR ratio is low (around 0.6) in the morning and evening but increases to around (1.3-1.4) at noon (Demotes-Mainard et al. 2016), which is consistent with our results. However, our observations showed that the daily mean R/FR ratio varies between 1.2 and 1.4 depending on the season, as it increased in summer and decreased in winter (Fig. 9). The decrease in the R/FR ratio in winter was significant (*p* < 0.001, *F* = 56.19), and the *ad* was approximately 13%. This may affect plant morphogenesis (Demotes-Mainard et al. 2016) and photosynthetic processes (Yang et al. 2020). Furthermore, the lower correlation between the R/FR ratio and DF suggests that the R/FR ratio is less dependent on weather conditions compared to other CWRs. Therefore, it is likely that the R/FR ratio functions as a signal for plants to perceive the seasons.

VP, which represents the water vapor in the atmosphere, was surprisingly not correlated with DF; this suggests that cloud cover may behave independently of VP, contrary to expectations. The UV-A/UV-B ratio correlated with VP, which could be evidence of an effect on plant UV signal perception. The quadratic model explained 61% of the total variation in the UV-A/UV-B ratio. The PAR/GSI ratio was also partially correlated to the VP, because the cubic model explained approximately 57% of the variation in the PAR/GSI ratio. In addition, the VP was strongly correlated with the R/FR ratio, and the linear model explained more than 77% of the total variation in the R/FR ratio. Therefore, the R/FR signal could be affected by the VP in the atmosphere and it may act as a signal for plants to perceive the VP.

The UV-A/UV-B ratio exhibited a significant correlation with AM, but only a weak correlation with DF. Previous research has demonstrated that while both UV-A and UV-B have a negative linear correlation with AM, UV-B has a steeper attenuation angle than does UV-A, which supports our findings (Borchi et al., 2011). The results suggest that AM may have an impact on UV-A/UV-B light signals in plants; this impact may be due to Rayleigh scattering, which primarily affects the short wavelength region. Field observations also provide evidence of the effect of AM on UV light (Riordan et al. 1990). The correlation between R/FR and PAR/GSI with AM was moderate (*r* < −0.5), and the developed model partially accounted for the variation, which reflected the combined effects of AM and other factors. Although some CWRs were correlated with AM, most correlations were weak (*r* < 0.4), and the developed model could not fully explain the variation (*R*^2^_adj_ < 0.1). Therefore, it appears that AM may only impact certain CWRs.

Previous research has shown that clouds can have varying effects at different wavelengths (Bartlett et al., 1998). However, Bartlett et al. (1998) argue that previous describing the effect of clouds on irradiance is inadequate due to the changing effect on the existence of different types of clouds. This study considered the DF to address this issue. We found that all CWRs, except for the UV-A/UV-B and R/FR ratios, had a strong relationship with the DF. Previous studies have shown that the UV-B/GSI ratio may increase under all-sky conditions unlike clear-sky conditions, which is consistent with our results (El-Nouby Adam and Ahmed, 2016). Our results demonstrate the significant impact of DF on plant light signaling. Plant light signals are crucial for controlling various physiological and metabolic processes in plants. Therefore, our findings highlight the significance of using DF in SR modeling and clustering, because it strongly influences plant light signals in addition to DF’s importance in canopy light distribution.

Furthermore, the multiple linear models showed that some of the environmental parameters, such as VP, AM, and DF, combined to influence specific light signals. The UV-A/UV-B light signal was best explained by the combined effects of VP and AM, whereas the R/FR and PAR/GSI light signals were best explained by the combined effects of VP and DF. This reveals the important interaction of parameters in changing plant light signals. Previous research supports our findings that VP affects the FR region of SR (Doroszewski et al. 2015). Our study expands on this knowledge by examining the seasonal variation in plant light signals and their relationship with VP, AM, and DF.

## Conclusions

Ground-based observations using high-precision spectroradiometers confirmed significant annual variations in the DF and all the CWRs considered. The high DFs during all seasons suggest that most of the days were cloudy or partly cloudy, indicating the limited applicability of clear-sky models in the Fukuoka region in Japan. Despite the higher VP in summer, the seasonal DF distribution was similar in summer and winter, whereas autumn showed a different DF distribution. The UV-A/UV-B, R/B, and R/G ratios increased in winter and decreased in summer. In contrast, the PAR/GSI, UV/GSI, UV/PAR, B/G, R/FR, and UV-B/B ratios increased in summer and decreased in winter. The monthly average PAR/GSI_(300-1100)_ ratio fluctuated significantly (0.43–0.47), which indicated that using a single multiplier to calculate GPP and plant growth modeling could lead to inaccurate calculations. The research found that AM and VP may influence the UV-A/UV-B and UV-B/B light signals and that VP is correlated with PAR/GSI and R/FR ratios. Additionally, the results revealed that DF may influence plant light signals and identified sources of variation in the CWRs, including the R/B, R/G, B/G, R/FR, and UV-B/B ratios other than UV ratios (UV/PAR and UV/GSI), except for the UV-A/UV-B ratio. The combined influence of VP and AM affects the UV-A/UV-B ratio, whereas the combined influence of VP and DF affects the R/FR and PAR/GSI light signals, and the combined influence of AM, VP, and DF affects the UV-B/B light signal. This research explains how VP, AM, and DF, individually and collectively affect plant light signals. These findings have implications for future research on plant ecophysiology and agricultural activities involved in plant photoreception.

## Abbreviations

SR, Solar radiation; UVR, ultraviolet radiation; PAR, photosynthetically active radiation; IR, infrared radiation; GPP, gross primary productivity; SSI, spectral solar irradiance; R/FR, red to far-red; R/B, red to blue; R/G, red to green; B/G, blue to green; UV-A/UV-B, DFR, diffuse solar radiation; DIR, direct solar radiation; GSI, global solar radiation; DF, diffuse factor/ diffuse ratio; LUE, light use efficiency; CWR, critical wavelength ratio; PPFD, photosynthetically active photon flux density; *rd,* relative difference; *ad,* absolute difference; VP, water vapor pressure; AM, airmass;

## Supporting information

Supplemental Figuers and Tables

## Acknowledgment

Tomoko Kawaguchi Akitsu provided technical advice on the spectroradiometer used in this study. The Faculty of Agriculture at Kyushu University supported the installation of the spectroradiometer. This work was also supported by the Japan Society for the Promotion of Science, KAKENHI Grant Number JP18H02511.

## Data availability

Data will be made available on request.

## Credit authorship contribution statement

Amila Nuwan Siriwardana: Formal analysis, Data curation, writing – original draft, Visualization. Atsushi Kume: Conceptualization, methodology, investigation, writing – review and editing, Funding acquisition.

## References

1. Akitsu TK, Nasahara KN, Ijima O, Hirose Y, Ide R, Takagi K, Kume A. 2022. The variability and seasonality in the ratio of photosynthetically active radiation to solar radiation: A simple empirical model of the ratio. International Journal of Applied Earth Observation and Geoinformation. 108:102724. 10.1016/J.JAG.2022.102724

2. Blal M, Khelifi S, Dabou R, Sahouane N, Slimani A, Rouabhia A, Ziane A, Neçaibia A, Bouraiou A, Tidjar B. 2020. A prediction models for estimating global solar radiation and evaluation meteorological effect on solar radiation potential under several weather conditions at the surface of Adrar environment. Measurement. 152:107348. 10.1016/J.MEASUREMENT.2019.107348

3. Borchi E, MacIi R, Bruzzi M, Scaringella M. 2011. Characterisation of SiC photo-detectors for solar UV radiation monitoring. Nucl Instrum Methods Phys Res A. 658(1):121–124. 10.1016/J.NIMA.2011.05.078

4. Brodersen CR, Vogelmann TC, Williams WE, Gorton HL. 2008. A new paradigm in leaf-level photosynthesis: direct and diffuse lights are not equal. Plant Cell Environ. 31(1):159–164. 10.1111/J.1365-3040.2007.01751.X

5. Chen M, Chory J, Fankhauser C. 2004. Light Signal Transduction in Higher Plants. 10.1146/annurev.genet38072902092259. 38:87–117. 10.1146/ANNUREV.GENET.38.072902.092259

6. Demotes-Mainard S, Péron T, Corot A, Bertheloot J, Le Gourrierec J, Pelleschi-Travier S, Crespel L, Morel P, Huché-Thélier L, Boumaza R, et al. 2016. Plant responses to red and far-red lights, applications in horticulture. Environ Exp Bot. 121:4–21. 10.1016/J.ENVEXPBOT.2015.05.010

7. Doroszewski A, Górski T, Kozyra J. 2015. Atmospheric moisture controls far-red irradiation: a probable impact on the phytochrome**. Int Agrophys. 29:283–289. 10.1515/intag-2015-0033

8. Durand M, Murchie EH, Lindfors A V., Urban O, Aphalo PJ, Robson TM. 2021. Diffuse solar radiation and canopy photosynthesis in a changing environment. Agric For Meteorol. 311:108684. 10.1016/J.AGRFORMET.2021.108684

9. Engerer NA, Mills FP. 2015. Validating nine clear sky radiation models in Australia. Solar Energy. 120:9–24. 10.1016/J.SOLENER.2015.06.044

10. Guechi A, Chegaar M, Aillerie M. 2013. Air Mass Effect on the Performance of Organic Solar Cells. Energy Procedia. 36:714–721. 10.1016/J.EGYPRO.2013.07.083

11. Kong SG, Okajima K. 2016. Diverse photoreceptors and light responses in plants. J Plant Res. 129(2):111–114. 10.1007/S10265-016-0792-5/FIGURES/1

12. Kono M, Noguchi K, Terashima I. 2014. Roles of the Cyclic Electron Flow Around PSI (CEF-PSI) and O 2-Dependent Alternative Pathways in Regulation of the Photosynthetic Electron Flow in Short-Term Fluctuating Light in Arabidopsis thaliana. Plant Cell Physiol. 55(5):990–1004. 10.1093/pcp/pcu033

13. Li D, Ju W, Lu D, Zhou Y, Wang H. 2015. Impact of estimated solar radiation on gross primary productivity simulation in subtropical plantation in southeast China. Solar Energy. 120:175–186. 10.1016/J.SOLENER.2015.07.033

14. Li T, Yang Q. 2015. Advantages of diffuse light for horticultural production and perspectives for further research. Front Plant Sci. 6(september):158826. 10.3389/FPLS.2015.00704/BIBTEX

15. Liu J, van Iersel MW. 2021. Photosynthetic Physiology of Blue, Green, and Red Light: Light Intensity Effects and Underlying Mechanisms. Front Plant Sci. 12:619987. 10.3389/FPLS.2021.619987/BIBTEX

16. Liu P, Tong X, Zhang Jinsong, Meng P, Li J, Zhang Jingru, Zhou Y. 2022. Effect of diffuse fraction on gross primary productivity and light use efficiency in a warm-temperate mixed plantation. Front Plant Sci. 13:966125. 10.3389/FPLS.2022.966125/BIBTEX

17. Lozano IL, Sánchez-Hernández G, Guerrero-Rascado JL, Alados I, Foyo-Moreno I. 2022. Analysis of cloud effects on long-term global and diffuse photosynthetically active radiation at a Mediterranean site. Atmos Res. 268:106010. 10.1016/J.ATMOSRES.2021.106010

18. Malitson HH. 1968. The solar electromagnetic radiation environment. Solar Energy. 12(2):197–203. 10.1016/0038-092X(68)90005-4

19. Mao J, Zhang YC, Sang Y, Li QH, Yang HQ. 2005. A role for Arabidopsis cryptochromes and COP1 in the regulation of stomatal opening. Proc Natl Acad Sci U S A. 102(34):12270– 12275. 10.1073/PNAS.0501011102/SUPPL_FILE/01011FIG7.PDF

20. Matthews JSA, Vialet-Chabrand S, Lawson T. 2020. Role of blue and red light in stomatal dynamic behaviour. J Exp Bot. 71(7):2253–2269. 10.1093/JXB/ERZ563

21. Mazza CA, Zavala J, Scopel AL, Ballaré CL. 1999. Perception of solar UVB radiation by phytophagous insects: Behavioral responses and ecosystem implications. Proc Natl Acad Sci U S A. 96(3):980. 10.1073/PNAS.96.3.980

22. Neugart S, Schreiner M. 2018. UVB and UVA as eustressors in horticultural and agricultural crops. Sci Hortic. 234:370–381. 10.1016/J.SCIENTA.2018.02.021

23. Orte F, Lusi A, Carmona F, D’Elia R, Faraminan A, Wolfram E. 2021. Comparison of NASA-POWER solar radiation data with ground-based measurements in the south of South America. 2021 19th Workshop on Information Processing and Control, RPIC 2021. 10.1109/RPIC53795.2021.9648428

24. Park Y, Runkle ES. 2023. Spectral-conversion film potential for greenhouses: Utility of green-to-red photons conversion and far-red filtration for plant growth. PLoS One. 18(2):e0281996. 10.1371/JOURNAL.PONE.0281996

25. Pennisi G, Blasioli S, Cellini A, Maia L, Crepaldi A, Braschi I, Spinelli F, Nicola S, Fernandez JA, Stanghellini C, et al. 2019. Unraveling the Role of Red:Blue LED Lights on Resource Use Efficiency and Nutritional Properties of Indoor Grown Sweet Basil. Front Plant Sci. 10. 10.3389/FPLS.2019.00305

26. Pennisi G, Orsini F, Blasioli S, Cellini A, Crepaldi A, Braschi I, Spinelli F, Nicola S, Fernandez JA, Stanghellini C, et al. 2019. Resource use efficiency of indoor lettuce (Lactuca sativa L.) cultivation as affected by red:blue ratio provided by LED lighting. Sci Rep. 9(1). 10.1038/S41598-019-50783-Z

27. Peratikou S, Charalambides AG. 2022. Estimating clear-sky PV electricity production without exogenous data. Solar Energy Advances. 2:100015. 10.1016/J.SEJA.2022.100015

28. Perliski LM, Solomon S. 1993. On the evaluation of air mass factors for atmospheric near-ultraviolet and visible absorption spectroscopy. Journal of Geophysical Research: Atmospheres. 98(D6):10363–10374. 10.1029/93JD00465

29. Polefka TG, Meyer TA, Agin PP, Bianchini RJ. 2012. Effects of Solar Radiation on the Skin. J Cosmet Dermatol. 11(2):134–143. 10.1111/J.1473-2165.2012.00614.X

30. Rai N, Morales LO, Aphalo PJ. 2021. Perception of solar UV radiation by plants: photoreceptors and mechanisms. Plant Physiol. 186(3):1382–1396. 10.1093/PLPHYS/KIAB162

31. Riordan CJ, Hulstrom RL, Myers DR. 1990. Influences of atmospheric conditions and air mass on the ratio of ultraviolet to total solar radiation. 10.2172/6344084

32. Sellaro R, Crepy M, Trupkin SA, Karayekov E, Buchovsky AS, Rossi C, Casal JJ. 2010a. Cryptochrome as a Sensor of the Blue/Green Ratio of Natural Radiation in Arabidopsis. Plant Physiol. 154(1):401–409. 10.1104/PP.110.160820

33. Sellaro R, Crepy M, Trupkin SA, Karayekov E, Buchovsky AS, Rossi C, Casal JJ. 2010b. Cryptochrome as a Sensor of the Blue/Green Ratio of Natural Radiation in Arabidopsis. Plant Physiol. 154(1):401–409. 10.1104/PP.110.160820

34. Shi Y, Ke X, Yang X, Liu Y, Hou X. 2022. Plants response to light stress. Journal of Genetics and Genomics. 49(8):735–747. 10.1016/J.JGG.2022.04.017

35. Shibuya K, Onodera S, Hori M. 2018. Toxic wavelength of blue light changes as insects grow. PLoS One. 13(6):e0199266. 10.1371/JOURNAL.PONE.0199266

36. Shinomura T, Nagatani A, Chory J, Furuya M. 1994. The Induction of Seed Germination in Arabidopsis thaliana Is Regulated Principally by Phytochrome B and Secondarily by Phytochrome A. Plant Physiol. 104(2):363–371. 10.1104/PP.104.2.363

37. Solanki SK, Krivova NA, Haigh JD. 2013. Solar Irradiance Variability and Climate. 10.1146/annurev-astro-082812-141007. 51:311–351. 10.1146/ANNUREV-ASTRO-082812-141007

38. Tan T, Li S, Fan Y, Wang Z, Ali Raza M, Shafiq I, Wang B, Wu X, Yong T, Wang X, et al. 2022. Far-red light: A regulator of plant morphology and photosynthetic capacity. Crop J. 10(2):300–309. 10.1016/J.CJ.2021.06.007

39. Tissot N, Ulm R. 2020. Cryptochrome-mediated blue-light signalling modulates UVR8 photoreceptor activity and contributes to UV-B tolerance in Arabidopsis. Nature Communications 2020 11:1. 11(1):1–10. 10.1038/s41467-020-15133-y

40. Trojak M, Skowron E, Sobala T, Kocurek M, Pałyga J. 2022. Effects of partial replacement of red by green light in the growth spectrum on photomorphogenesis and photosynthesis in tomato plants. Photosynth Res. 151(3):295–312. 10.1007/S11120-021-00879-3/FIGURES/6

41. Earthdata. earthdata.nasa.gov [accessed 2024 May 5]. https://www.earthdata.nasa.gov/topics/atmosphere/atmospheric-water-vapor/water-vapor-indicators/vapor-pressure

42. Wang B, Yue X, Zhou H, Zhu J. 2022. Impact of diffuse radiation on evapotranspiration and its coupling to carbon fluxes at global FLUXNET sites. Agric For Meteorol. 322:109006. 10.1016/J.AGRFORMET.2022.109006

43. Wang S, Liu Xiaoting, Liu Xiaoning, Xue J, Ren X, Zhai Y, Zhang X. 2022. The red/blue light ratios from light-emitting diodes affect growth and flower quality of Hippeastrum hybridum ‘Red Lion.’ Front Plant Sci. 13:1048770. 10.3389/FPLS.2022.1048770/BIBTEX

44. Wu J, Fang H, Qin W, Wang L, Song Y, Su X, Zhang Y. 2022. Constructing High-Resolution (10 km) Daily Diffuse Solar Radiation Dataset across China during 1982–2020 through Ensemble Model. Remote Sens (Basel). 14(15). 10.3390/RS14153695

45. Xin Q, Gong P, Suyker AE, Si Y. 2016. Effects of the partitioning of diffuse and direct solar radiation on satellite-based modeling of crop gross primary production. International Journal of Applied Earth Observation and Geoinformation. 50:51–63. 10.1016/J.JAG.2016.03.002

46. Yadav A, Singh D, Lingwan M, Yadukrishnan P, Masakapalli SK, Datta S. 2020. Light signaling and UV-B-mediated plant growth regulation. J Integr Plant Biol. 62(9):1270– 1292. 10.1111/JIPB.12932

47. Yang J, Li C, Kong D, Guo F, Wei H. 2020. Light-Mediated Signaling and Metabolic Changes Coordinate Stomatal Opening and Closure. Front Plant Sci. 11:601478. 10.3389/FPLS.2020.601478/BIBTEX

48. Yoshino MM, Kazuko U. 1981. Regionality of climatic change in East Asia. GeoJournal. 5(2):123–132. 10.1007/BF02582045/METRICS

49. Yuan W, Cai W, Xia J, Chen J, Liu S, Dong W, Merbold L, Law B, Arain A, Beringer J, et al. 2014. Global comparison of light use efficiency models for simulating terrestrial vegetation gross primary production based on the LaThuile database. Agric For Meteorol. 192–193:108–120. 10.1016/J.AGRFORMET.2014.03.007

