## Supplemental Figuers and Tables for "Annual Variability in Diffuse Ratio and Spectral Characteristics of Solar Radiation: Cloud Effects in a Temperate Monsoon Region"

**Supplementary legends**

**Sup. 1** Meteorological information for the site **(a)** Daily mean temperature by month **(b)** Daily mean humidity by month **(c)** Daily mean local air pressure by month **(d)** Annual variation in actual and saturated vapor pressure. **(e)** Daily mean AM by month **(f)** Daytime $GSI_{(200\text{-}1200)}$ quantum flux vs. all-day $GSI_{(actual)}$ quantum flux, during 2021

**Sup. 2** Comparative analysis of CWRs and DF across seasons: This table presents the results of a Pearson's correlation test, examining the distribution of percentage days across different seasons for DF and various CWRs. Pearson's correlation coefficients and level of significance provide a quantitative measure of the similarity in distributions of DF and CWRs between seasons.

**Sup. 3** Seasonal subplot of the percentage of days within each DF range., The percentage of days distribution between different DF ranges among seasons was plotted for the same DF percentile range as Fig.1b, to observe the distribution similarity between different seasons during 2021.

**Sup. 4** Daily mean GSI vs. daily mean DF, and the linear model fitted to approximate the variation in GSI during 2021.

**Sup. 5** Annual variation of UV/GSI photon flux ratios, raincloud plot with box plot constructed to observe the variations by month.

**Sup. 6** Comparative seasonal distribution of days within various UV ratio ranges. This figure presents a detailed analysis of the parentage distribution of days within different UV ratio percentiles across various seasons. The same UV ratio ranges were used to create seasonal subplots for comparative observation of seasonal variations. **(a)** seasonal distribution of days within different ranges of UV-B/B ratio. **(b)** seasonal distribution of days within different ranges of UV/PAR ratio. **(c)** seasonal distribution of days within different ranges of UV-A/UV-B quantum flux ratio during 2021.

**Sup. 7** Multiple comparison test results for UV ratios. **(a)** Multiple test results for the UV-B/B ratio, **(b)** multiple test results for the UV/PAR ratio, and **(c)** multiple test results for the UV-A/UV-B ratio

**Sup. 8** Comparative seasonal distribution of days within various R, B and G ratio ranges. This figure presents a detailed analysis of the parentage distribution of days within different R, B and G ratio percentiles across various seasons. The same ratio ranges were used to create seasonal subplots for each R, B and G ratio to comparative observation of seasonal variations., **(a)** Seasonal distribution of dates in different ranges of R/B photon flux ratio. **(b)** Seasonal distribution of dates in different ranges of R/G photon flux ratio. **(c)** Seasonal distribution of dates in different ranges of B/G photon flux ratio during 2021.

**Sup. 9** Multiple comparison test results for R, B, and G ratios. **(a)** Multiple test results for the R/B ratio, **(b)** multiple test results for the R/G ratio, and **(c)** multiple test results for the B/G ratio.

**Sup. 10** Comparative seasonal distribution of days within various R/FR ratio ranges. The figure presents a detailed analysis of the parentage distribution of days within different R/FR ratio percentiles across various seasons. The same ratio range was used to create seasonal subplots for comparative observation of seasonal variations of dates in different ranges of R/FR photon flux ratio during 2021.

Sup. 1

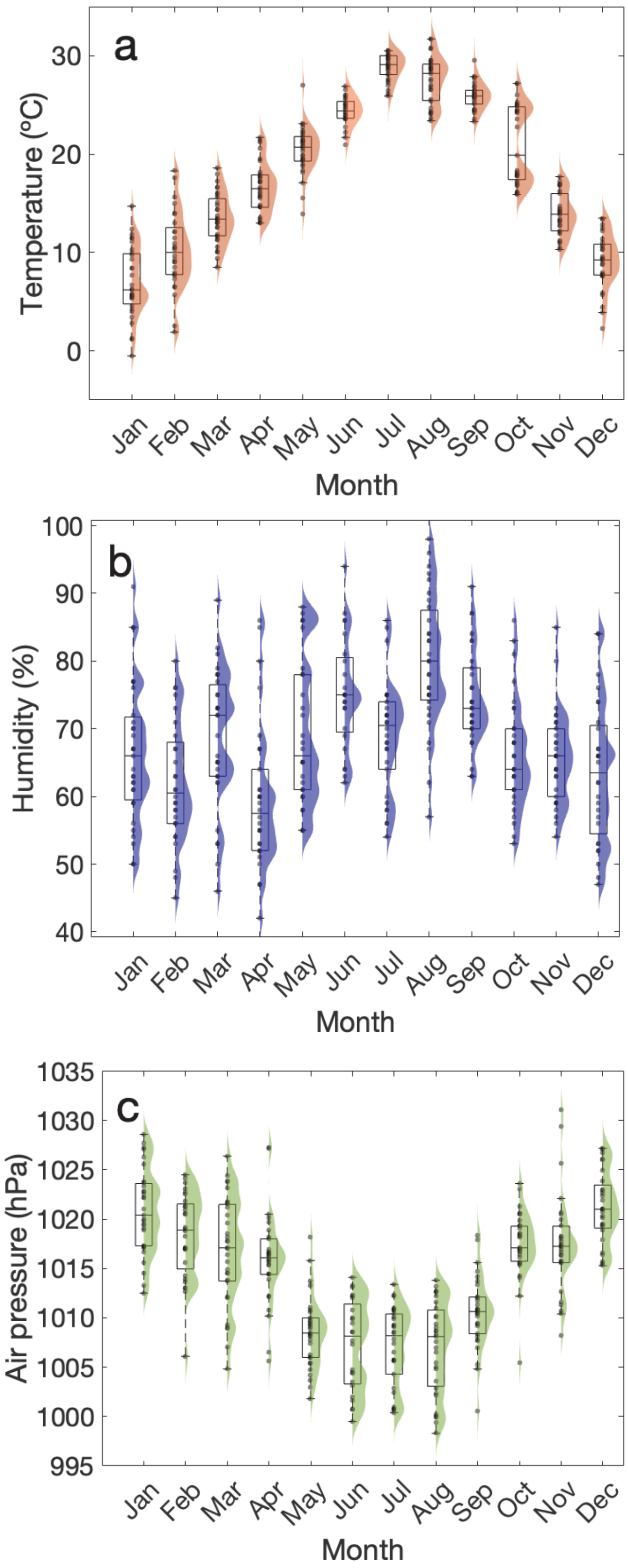
a
Temperature (°C)
30
20
10
0
Jan
Feb
Mar
Apr
May
Jun
Jul
Aug
Sep
Oct
Nov
Dec
Month
b
Humidity (%)
100
90
80
70
60
50
40
Jan
Feb
Mar
Apr
May
Jun
Jul
Aug
Sep
Oct
Nov
Dec
Month
c
Air pressure (hPa)
1035
1030
1025
1020
1015
1010
1005
1000
995
Jan
Feb
Mar
Apr
May
Jun
Jul
Aug
Sep
Oct
Nov
Dec
Month

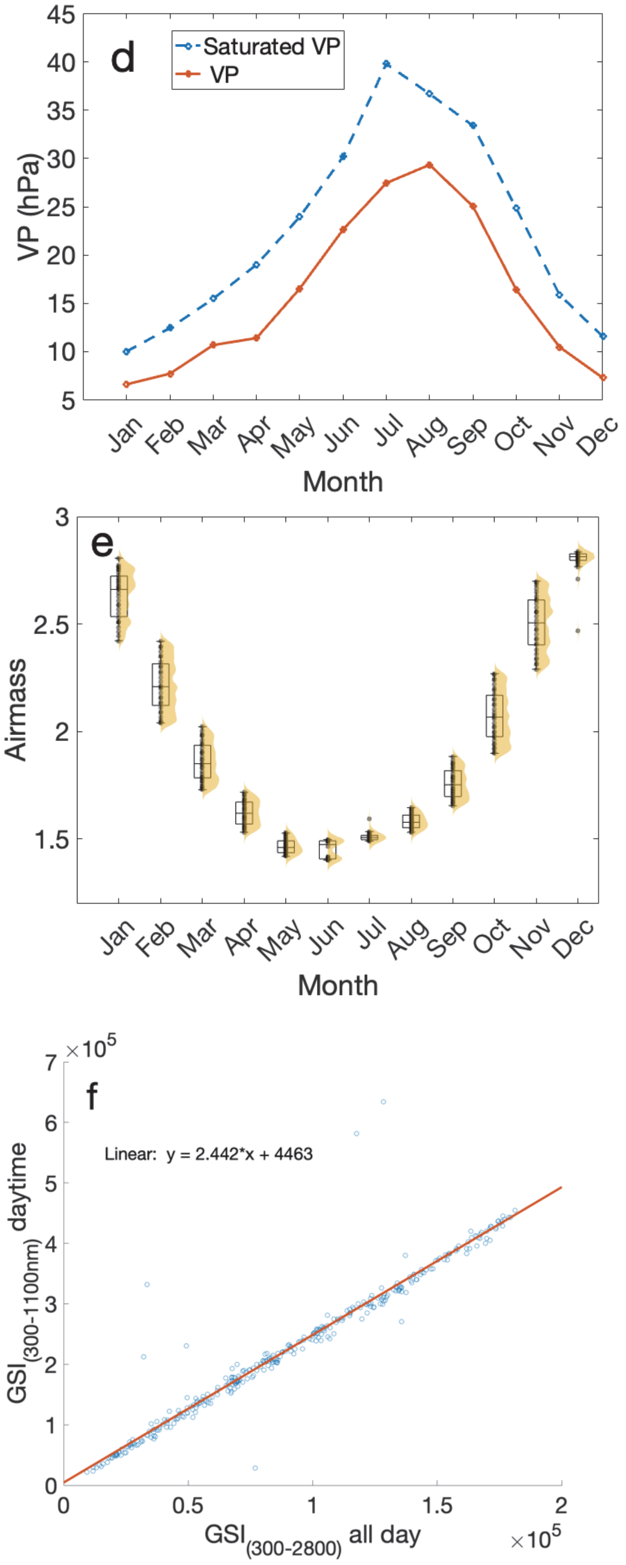
d
Saturated VP
VP
VP (hPa)
45
40
35
30
25
20
15
10
5
Jan
Feb
Mar
Apr
May
Jun
Jul
Aug
Sep
Oct
Nov
Dec
Month
e
Airmass
3
2.5
2
1.5
Jan
Feb
Mar
Apr
May
Jun
Jul
Aug
Sep
Oct
Nov
Dec
Month
f
$\times 10^5$
7
6
5
4
3
2
1
0
Linear: y = 2.442*x + 4463
$GSI_{(300-1100nm)}$ daytime
0
0.5
1
1.5
2
$GSI_{(300-2800)}$ all day
$\times 10^5$

**Sup. 2**

| | | **Seasons** | | | DR/CWRs correlation of % dates distribution between seasons | | | | | | | |
|---|---|---|---|---|---|---|---|---|---|---|---|---|
| | | **winter** | **spring** | **autumn** | DR | UV-B/B | UV/PAR | UV-A/UV-B | R/B | R/G | B/G | R/FR |
| **Seasons** | **Summer** | ✔ | | | 0.76 | 0.33 | 0.04 | 0.16 | 0.19 | 0.13 | $0.87^{*}$ | 0.05 |
| | **Summer** | | ✔ | | $0.95^{**}$ | 0.38 | 0.27 | 0.41 | 0.36 | 0.81 | $0.88^{*}$ | 0.01 |
| | **Summer** | | | ✔ | 0.40 | 0.09 | 0.30 | 0.37 | 0.33 | 0.5 | $0.94^{**}$ | 0.23 |
| | **winter** | | ✔ | | $0.78^{*}$ | 0.01 | 0.75 | 0.04 | $0.94^{**}$ | 0.52 | $0.96^{**}$ | 0.50 |
| | **winter** | | | ✔ | 0.10 | 0.27 | $0.83^{*}$ | 0.16 | $0.95^{**}$ | $0.83^{*}$ | $0.97^{**}$ | 0.31 |
| | **spring** | | | ✔ | 0.36 | 0.87 | $0.97^{**}$ | $0.91^{*}$ | $1.00^{***}$ | $0.88^{*}$ | $0.98^{**}$ | $0.84^{*}$ |

$^{***}p < 0.001$, $^{**}p < 0.01$, and $^{*}p < 0.05$

**Sup. 3**

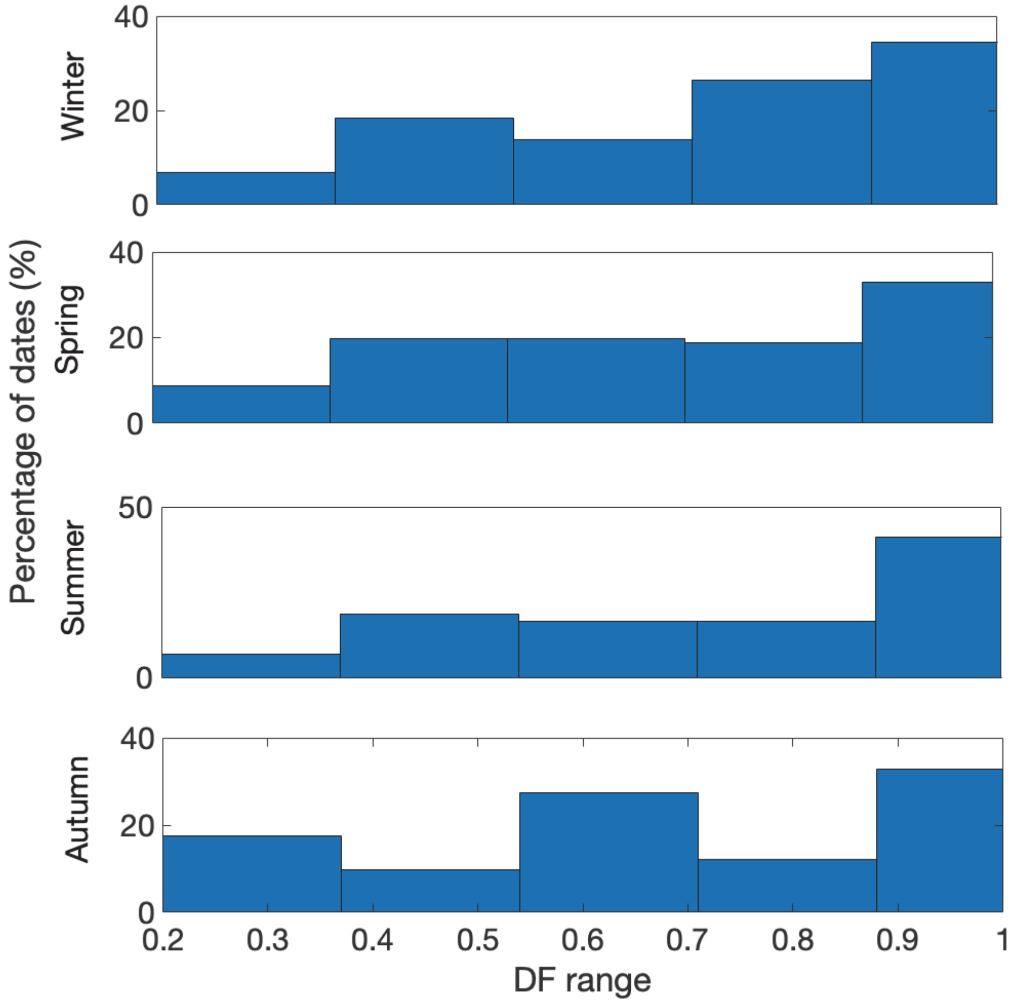

Percentage of dates (%)
Winter
Spring
Summer
Autumn
40
20
0
40
20
0
50
0
40
20
0
0.2
0.3
0.4
0.5
0.6
0.7
0.8
0.9
1
DF range

**Sup. 4**

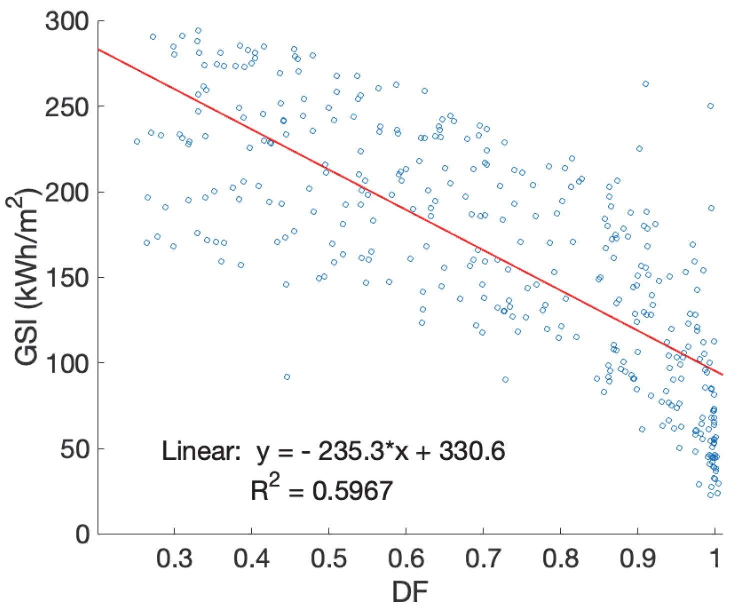

300
250
200
150
100
50
0
GSI (kWh/m²)
0.3
0.4
0.5
0.6
0.7
0.8
0.9
1
DF
Linear: y = - 235.3*x + 330.6
R² = 0.5967

**Sup. 5**

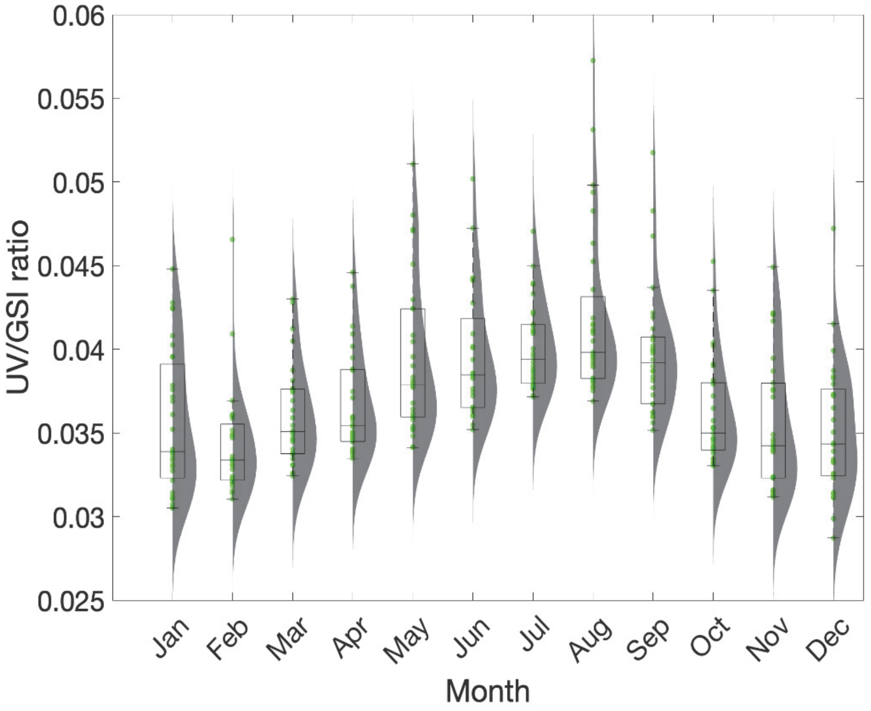
0.06
0.055
0.05
0.045
0.04
0.035
0.03
0.025
UV/GSI ratio
Jan
Feb
Mar
Apr
May
Jun
Jul
Aug
Sep
Oct
Nov
Dec
Month

**Sup 6**

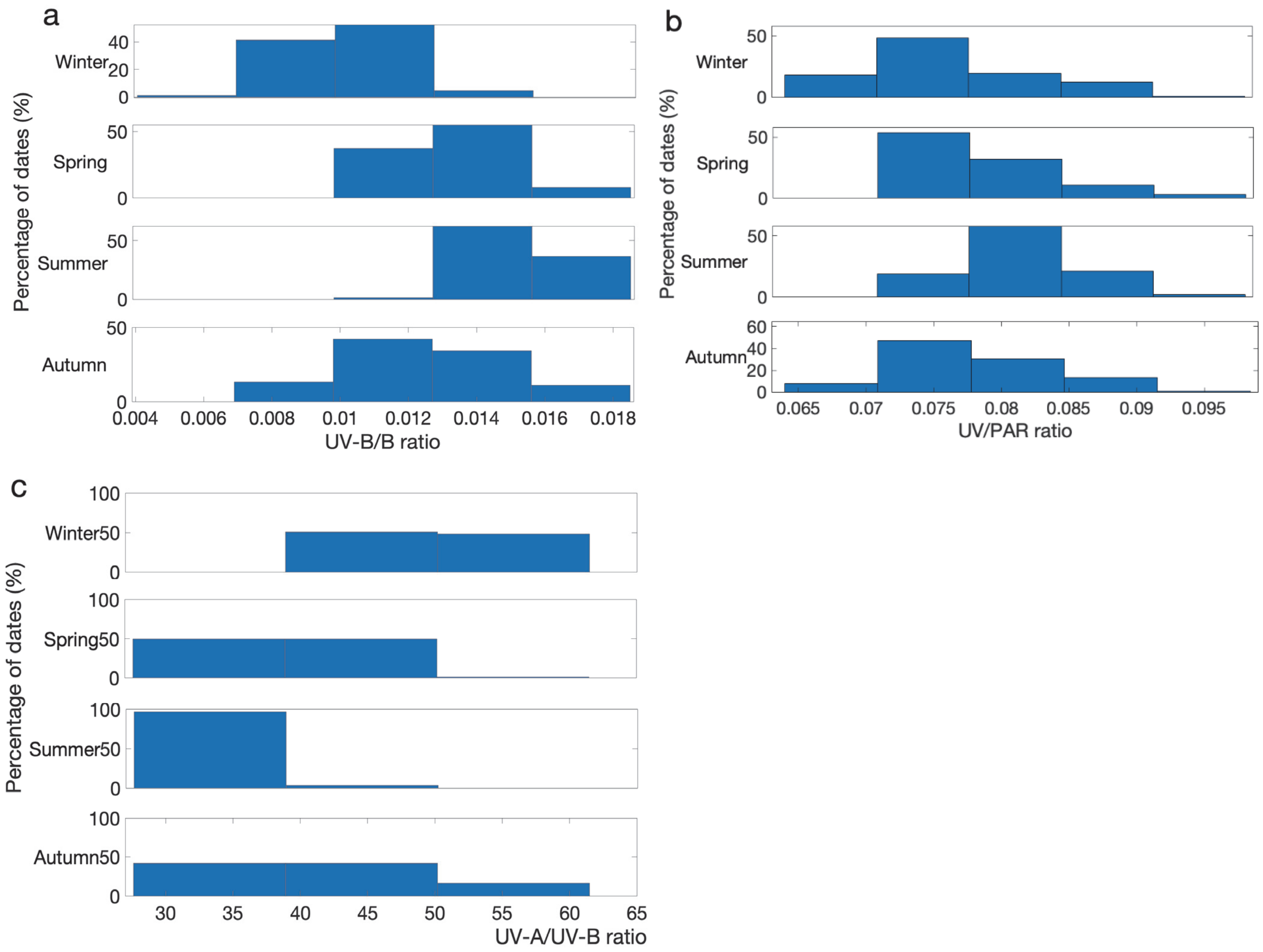
a
Winter
Spring
Summer
Autumn
Percentage of dates (%)
0.004 0.006 0.008 0.01 0.012 0.014 0.016 0.018
UV-B/B ratio
b
Winter
Spring
Summer
Autumn
Percentage of dates (%)
0.065 0.07 0.075 0.08 0.085 0.09 0.095
UV/PAR ratio
c
Winter
Spring
Summer
Autumn
Percentage of dates (%)
30 35 40 45 50 55 60 65
UV-A/UV-B ratio

**Sup. 7**

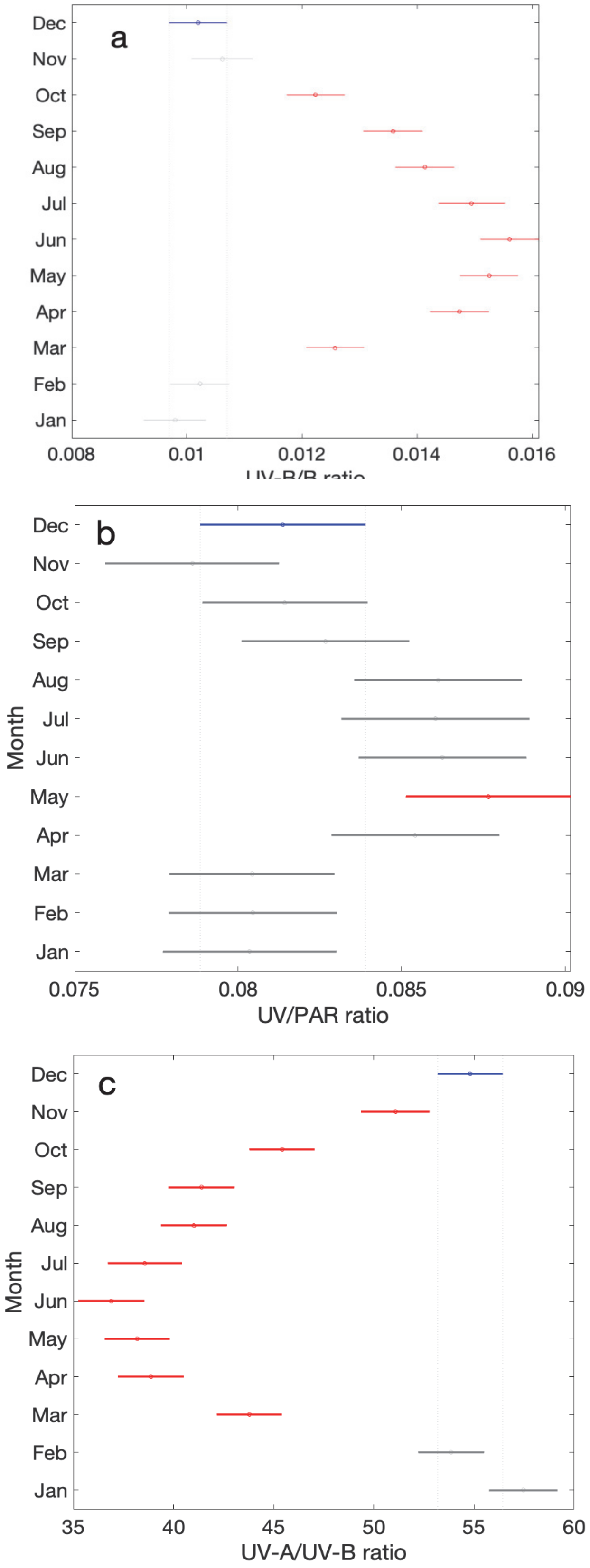
a
Dec
Nov
Oct
Sep
Aug
Jul
Jun
May
Apr
Mar
Feb
Jan
0.008
0.01
0.012
0.014
0.016
b
Month
0.075
0.08
0.085
0.09
UV/PAR ratio
c
35
40
45
50
55
60
UV-A/UV-B ratio

**Sup. 8**

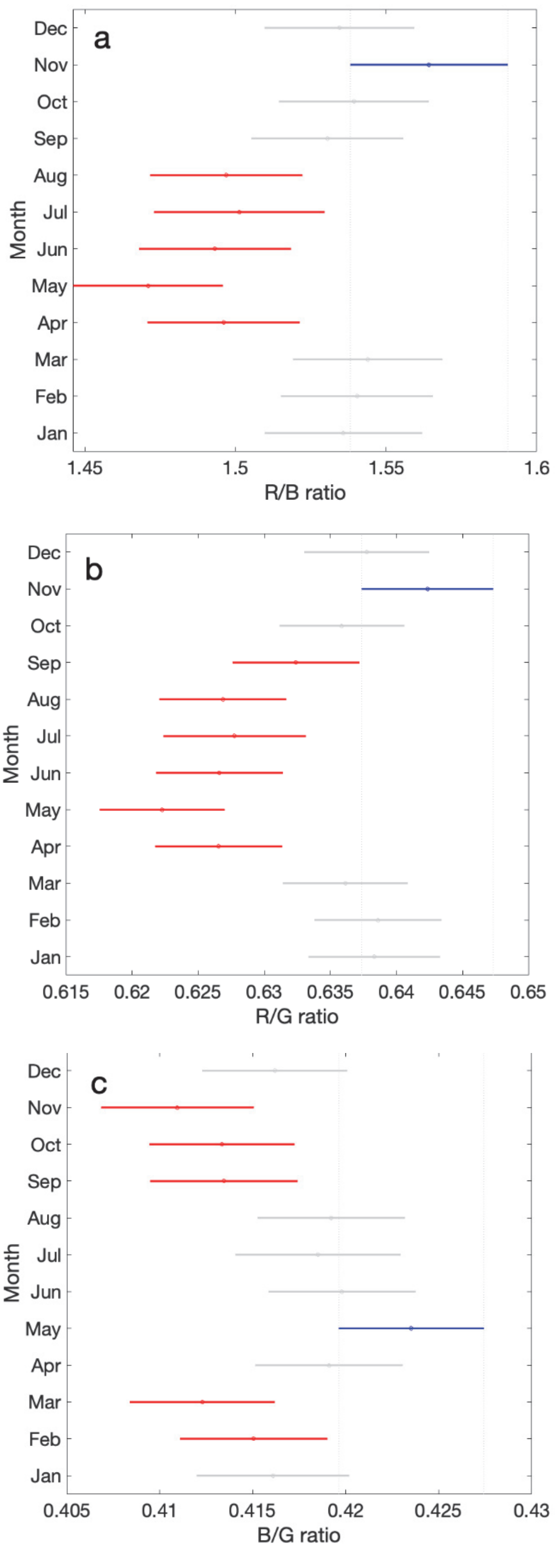
a
b
c
Month
Dec
Nov
Oct
Sep
Aug
Jul
Jun
May
Apr
Mar
Feb
Jan
1.45
1.5
1.55
1.6
R/B ratio
0.615
0.62
0.625
0.63
0.635
0.64
0.645
0.65
R/G ratio
0.405
0.41
0.415
0.42
0.425
0.43
B/G ratio

**Sup. 9**

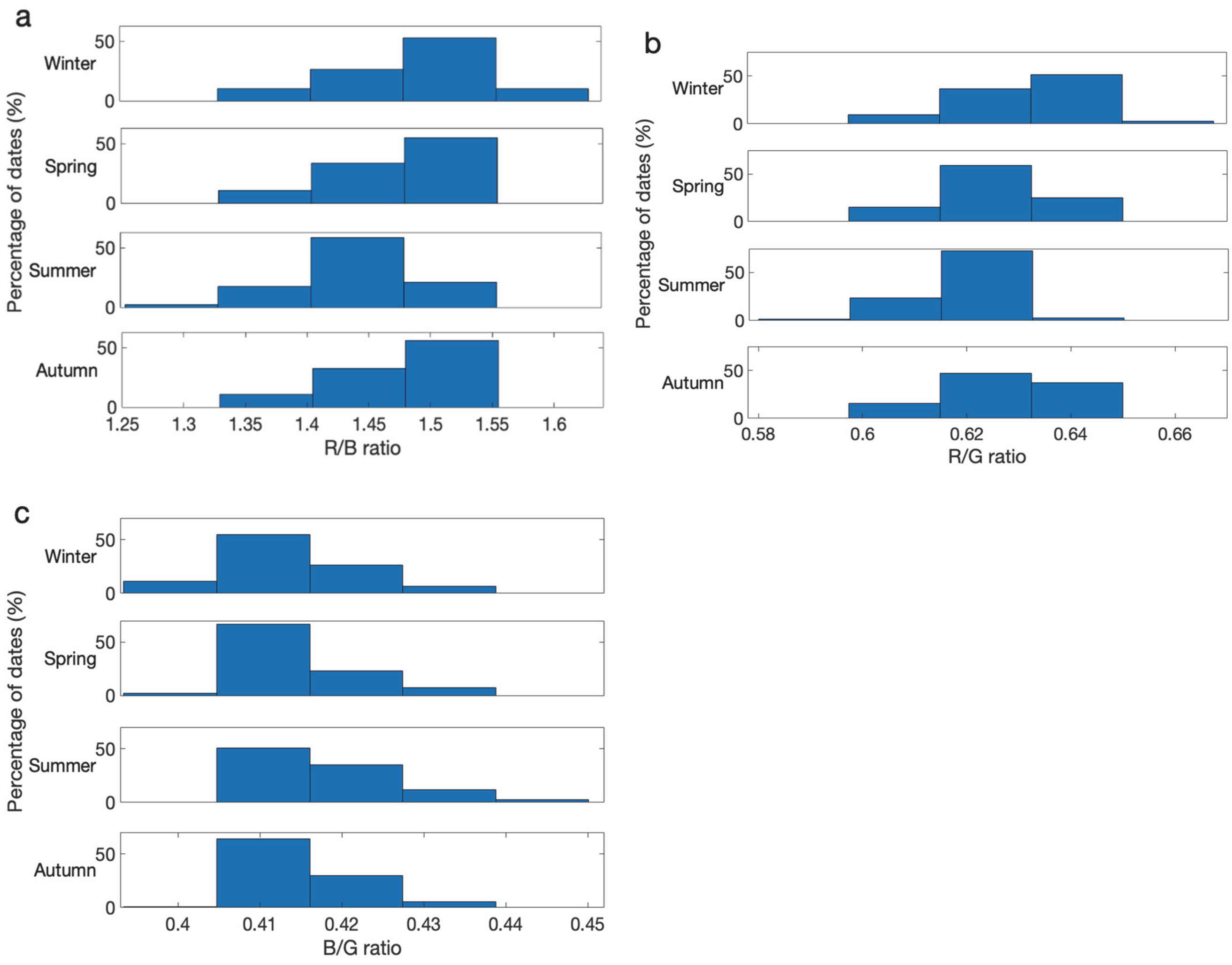

a
Winter
Spring
Summer
Autumn
Percentage of dates (%)
50
0
1.25
1.3
1.35
1.4
1.45
1.5
1.55
1.6
R/B ratio
b
R/G ratio
0.58
0.6
0.62
0.64
0.66
c
B/G ratio
0.4
0.41
0.42
0.43
0.44
0.45

**Sup. 10**

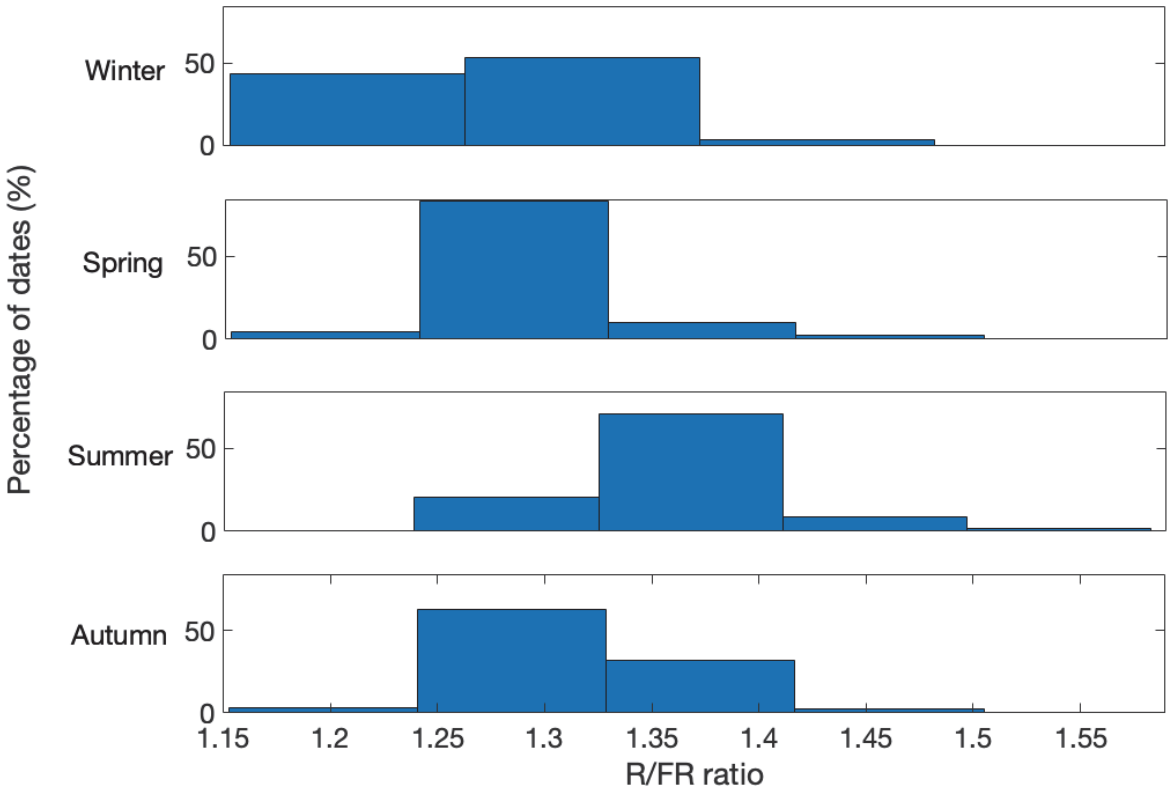

Percentage of dates (%)
Winter
50
0
Spring
50
0
Summer
50
0
Autumn
50
0
1.15
1.2
1.25
1.3
1.35
1.4
1.45
1.5
1.55
R/FR ratio
